# Neural and computational underpinnings of biased confidence in human reinforcement learning

**DOI:** 10.1101/2023.03.08.531656

**Authors:** Chih-Chung Ting, Nahuel Salem-Garcia, Stefano Palminteri, Jan B. Engelmann, Maël Lebreton

## Abstract

While navigating a fundamentally uncertain world, humans and animals constantly produce subjective confidence judgments, thereby evaluating the probability of their decisions, actions or statements being correct. Confidence typically correlates with neural activity positively in a ventromedial-prefrontal (VMPFC) network and negatively in a dorsolateral and dorsomedial prefrontal network. Here, combining fMRI with a reinforcement-learning paradigm, we leverage the fact that humans are more confident in their choices when seeking gains than avoiding losses to reveal a functional dissociation: whereas the dorsal prefrontal network correlates negatively with a condition-specific confidence signal, the VMPFC network positively encodes task-wide confidence signal incorporating the valence-induced bias. Challenging dominant neuro-computational models, we found that decision-related VMPFC activity better correlates with confidence than with option-values inferred from reinforcement-learning models. Altogether, these results identify the VMPFC as a key node in the neuro-computational architecture that builds global feeling-of-confidence signals from latent decision variables and contextual biases during reinforcement-learning.

## Introduction

Humans and animals seem to be constantly engaged in computing the subjective probability of having made the right choice, having successfully memorized or recognized a cue, having correctly executed the desired action or having endorsed the most truthful statement–, thereby producing automatic confidence judgments (Fleming & Daw, 2017; Fleming & Dolan, 2012; Lebreton et al., 2015; Pouget et al., 2016; Yeung & Summerfield, 2012). These metacognitive confidence judgments are increasingly considered as having a critical functional role in (sequential) decision-making, controlling the integration of new evidence (Desender et al., 2018), adjusting speed-accuracy trade-offs (van den Berg et al., 2016), and triggering changes of mind (Fleming et al., 2018; Folke et al., 2016). Likewise, a recent but increasing number of studies suggests that confidence could be a key variable to understand human (reinforcement-) learning behaviour both at the normative and descriptive levels (Boldt et al., 2019; Cortese et al., 2020; Hainguerlot et al., 2018; Heilbron & Meyniel, 2019; Meyniel, 2020; Vaghi et al., 2017).

At the neurobiological levels, the computation of confidence and the production of confidence judgments has been consistently associated with neural activity in two main prefrontal networks across a large variety of cognitive tasks: a negative prefrontal network, encompassing dorsal anterior cingulate cortex (dACC), bilateral insula, dorso-medial and dorsolateral prefrontal cortices, and a positive ventral network, mostly centered around the ventromedial prefrontal cortex (Cortese, 2021; Morales et al., 2018; Rouault, Lebreton, et al., 2022; Vaccaro & Fleming, 2018; White et al., 2014). For instance, dACC was originally identified as a key centre for performance monitoring and error detection (Holroyd & Coles, 2002; Taylor et al., 2007) as well as for the computation of uncertainty-related variables (Behrens et al., 2007), before being more generally integrated as a part of a large network negatively correlating with confidence judgments (Bang & Fleming, 2018; Boldt & Yeung, 2015; Heereman et al., 2015; Morales et al., 2018; Rouault, Lebreton, et al., 2022). More recently, BOLD activity in the ventromedial prefrontal cortex (VMPFC) and pregenual anterior cingulate cortex (pgACC) has been positively associated with confidence and self-performance evaluation, first in the context of value-based decision-making (De Martino et al., 2013), and then more broadly in other contexts and tasks (Bang & Fleming, 2018; Gherman & Philiastides, 2018; Hoven, et al., 2022; Lebreton et al., 2015; Morales et al., 2018; Rouault et al., 2022).

While both positive and negative prefrontal networks are omnipresent in the most-recent meta-analyses and theories of confidence and metacognition judgments (Cortese, 2021; Vaccaro & Fleming, 2018) there is, to date, very little empirical evidence to formally dissociate the relative roles of those two networks in the computation of confidence – but see e.g. (Bang & Fleming, 2018; Cortese, 2021). One promising hypothesis is that some of those network elements could be involved in different stages of confidence processing, including computing and integrating different confidence-building variables such as levels of uncertainty. Uncertainty and confidence can indeed be distinguished at the theoretical and computational levels: while confidence can be defined as the probability that a decision (or a proposition) is correct given the evidence, (un)certainty refers to the encoding of all other probability distributions over sensory and cognitive variables on which choices and confidence are ultimately built (Cortese, 2021; Fleming & Daw, 2017; Pouget et al., 2016). Thereby, these two quantities might be easily confoundable ‒potentially explaining why they have been associated with similar brain regions and neural patterns of activity in previous studies‒ but remain theoretically dissociable. Given the previous association of the negative network with uncertainty and error detection (Yeung & Summerfield, 2012), and of the positive network with affect and subjective valuation (Lieberman et al., 2019), one credible neurocomputational architecture would ascribe to the negative network a role in representing objective uncertainty –which often (negatively) correlates with confidence–, and to the VMPFC a role in aggregating a composite variable corresponding to the subjective, phenomenological feeling of confidence, from decision-related uncertainty variables and all other incidental signals influencing confidence.

Here, to test this putative architecture, we leverage a reinforcement learning paradigm that naturally orthogonalizes specific dimensions of difficulty and affective information (**Figure 1 A-B**), by factorially manipulating two features of choice outcomes: their valence (monetary gains or losses) and the quantity of information (partial versus complete feedback). Our idea is to take advantage of the valence-induced bias in confidence judgments described in the context of this task – i.e. the fact that participants are genuinely more confident in their choices when seeking gains than avoiding losses, despite identical objective difficulty and learning performance (Lebreton, et al., 2019; Salem-Garcia et al., 2021; Ting et al., 2020) – **Figure 1C**. Considering the task features and the typical participant behavior, a brain region encoding *objective uncertainty* should therefore correlate with confidence in all conditions, and exhibit signal differences between complete and partial-information contexts, as the objective uncertainty is higher in partial than complete-information contexts. On the other side of the spectrum, a brain region encoding *task-wide confidence* (corresponding to the reported, absolute feeling of confidence) should correlate with confidence in all conditions, and exhibit signal differences between gain and loss contexts, as participants report higher confidence in a gain context (despite similar choice difficulty and performance observed in a loss context). Finally, we also define a third variable, *condition-specific confidence*, which simply indexes the relative increase of confidence in each learning context due to the incremental improvement of choice accuracy caused by feedback-based learning. A brain region encoding condition-specific confidence should therefore correlate with confidence in all contexts, but not exhibit any signal difference due to our manipulation of valence and information (**Figure 1D)**.

**Figure 1.**
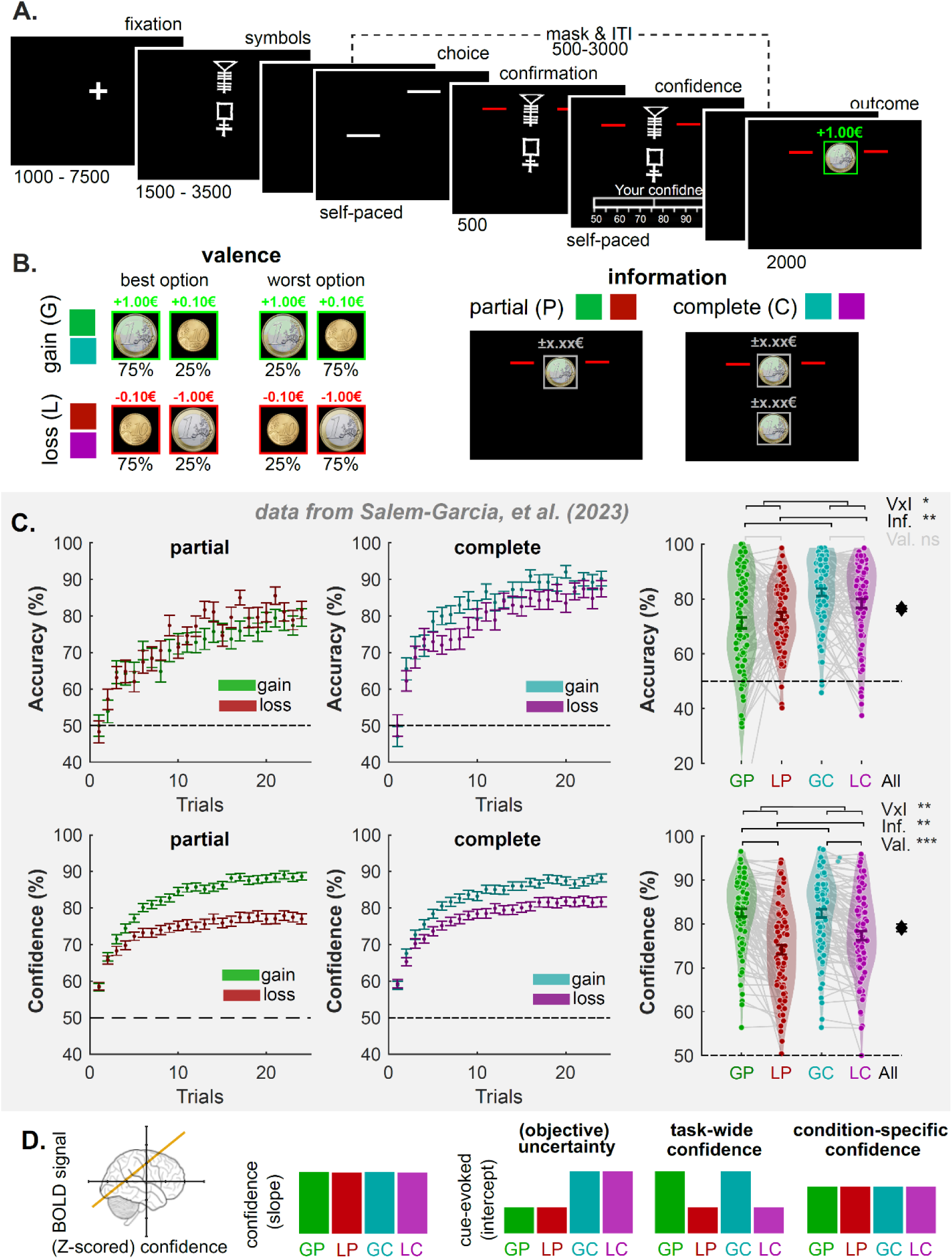
Experimental design and hypotheses. **(A)** Successive screens displayed during the learning task. Durations are given in ms. **(B)** Illustration of two-by-two factorial design with outcome valance (gain and loss) and information (partial and complete) manipulations. Each condition is consistently associated with a pair of symbols in each run. Each symbol is consistently associated with a probability (75% or 25%) of getting larger gains (€+1.0) and smaller gains (€+0.1) in the gain conditions and is consistently associated with a probability of getting smaller losses (€-0.1) and larger losses (€-1.0) in the loss conditions. In the outcome phase, the outcome from the chosen symbol is always displayed and highlighted with two red bars regardless of the information condition. The outcome from the unchosen option is absent in the partial information condition but is available in the complete information condition. **(C)** Evolution of average accuracy (upper panels) and confidence (bottom panels) across trials from five instrumental learning tasks (N = 90), which were reanalyzed and reported in Salem-Garcia et al., 2022. Different colors represent different contexts following the conventions from panel A. **(D)** Qualitative predictions about the relationship between brain activation patterns (BOLD signal) and confidence (e.g., the yellow line), for three possible confidence-related signals: uncertainty, condition-specific confidence, and task-wide confidence. The relationships can be summarized with a slope and an intercept (cue-evoked), across conditions. GP: gain/partial; LP: loss/partial; GC: gain/complete; LC: loss/complete; Val.: Valence manipulation; Inf. Information manipulation. V×I: Valence and information interaction. ∼: .05<*P*<.1; *: .001<*P*<.01; **: .01<*P*<.001; ***: *P*<.001

Following this reasoning, we recorded BOLD activity in participants while they performed the reinforcement-learning task featuring manipulations of outcome valence and information quality, paired with confidence elicitations. Behavioral analyses first confirmed the presence of the valence-induced confidence bias. fMRI analyses showed that the confidence was positively and negatively related to the activity in the prefrontal networks regardless of affective information and task difficulty manipulations. Using theory-driven qualitative patterns of activation as well as a quantitative model comparison exercise, our neuro-imaging analyses then revealed a functional dissociation. On the one hand, neural activity in the negative prefrontal network (i.e., DMPFC and DLPFC) correlated with a condition-specific confidence signal that gradually builds up, independently in each learning context. On the other hand, neural activity in the positive prefrontal network (i.e., VMPFC) additionally integrates contextual effects such as the valence-induced confidence bias, thereby representing absolute, task-wide confidence that mimics the feeling-of-confidence reported by participants. We further verified the role of the positive network in reinforcement learning via model-based fMRI analysis. In short, while VMPFC was also engaged in the computational process, the activity in the VMPFC can be better explained by confidence than other ongoing computational variables, including chosen option values and value differences.

## Results

Forty participants took part in our experiment and completed the instrumental learning task in the MRI scanner. During the learning task (**Figure 1A**), participants repeatedly faced pairs of abstract symbols (cues), that were probabilistically associated with monetary outcomes (gains or losses). In each pair, also referred to as context, one cue was associated with a better expected outcome (i.e., higher probability of gain or lower probability of loss), and the goal of participants was to learn, by trial and error, to identify and preferentially choose this cue. Two main contextual factors were orthogonally manipulated: outcome valence and outcome information (Lebreton, et al., 2019; Palminteri et al., 2015; Ting et al., 2020). The valence factor defines Gain and Loss contexts, which respectively only include cues probabilistically associated with gains or losses (**Figure 1B**). The information factor defines Partial and Complete information contexts, where feedback is respectively provided only for the chosen cue, or for both the chosen and unchosen cues (**Figure 1C**). In addition, at each trial, participants reported their confidence in their choice on a probabilistic scale as the subjective probability of having made a correct choice from 50% indicating chance level to 100% (indicating certainty). Those confidence judgments were incentivized using a matching probability mechanism –see **Methods** and (Hollard et al., 2016; Schlag et al., 2015) for details. Note that we decoupled the decision and response-related processes by delaying the mapping between the cue and the motor response, so as to minimize the inherent correlation between decision response times and confidence judgments –see **Figure 1A**, **Methods** and (Ting et al., 2020) for details.

## Reinforcement-learning behavior feature the valence-induced confidence bias

Overall, participants’ choice accuracy (i.e., the average probability of choosing the better symbol) is above guessing level (t_39_ = 17.78; *P* < .001; **Supplementary Table S1**), indicating that they were able to identify and select the better symbols from the probabilistic outcomes, by trial and error. We then evaluated the effects of our main experimental factors on the two behavioral variables of interest: choice accuracy and confidence judgments (**Figure 2**). Replicating previous reports (Fontanesi et al., 2019; Lebreton, et al., 2019; Palminteri et al., 2015; Salem-Garcia et al., 2021; Ting et al., 2020), we confirmed that choice accuracy is modulated by information but not valence (two-way repeated-measures ANOVA: valence F_1,39_ = 0.00, P = .9666; information F_1,39_ = 22.05, *P* <.001; interaction: F_1,39_ = 0.01, *P* = .9056).

**Figure 2.**
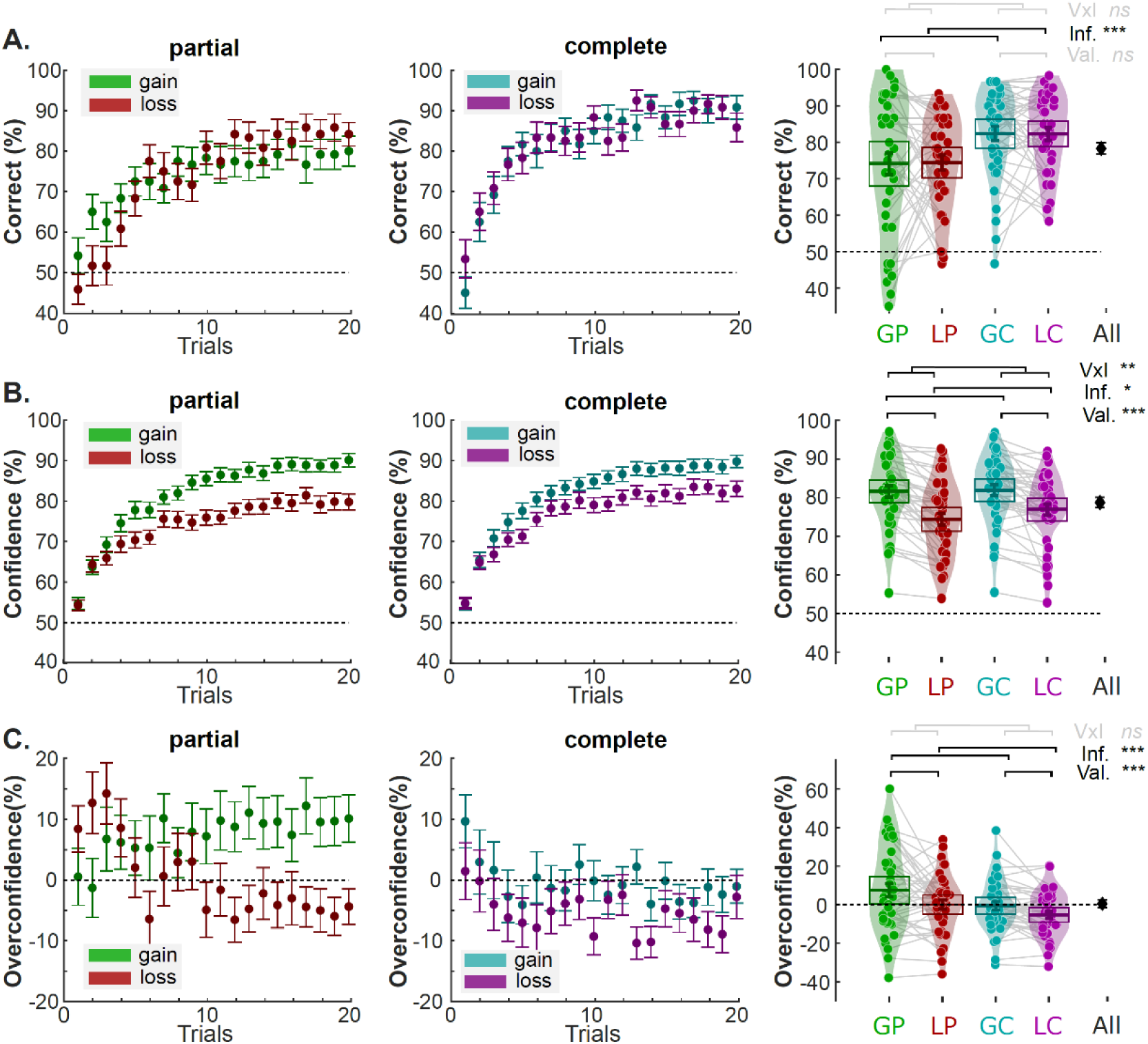
The effect of outcome valence and information on learning and confidence. Left and middle panels are trial-by-trial **(A)** percentage of correct responses, **(B)** Confidence rating, and **(C)** overconfidence in the partial information (left panels) and complete information condition (middle panels). Dots and error bars represent the trial-resolved mean ± SEM of the participant data. Right panels picture condition-specific averages. **(A)** percentage of correct responses, **(B)** confidence rating, and **(C)** overconfidence across conditions at the individual level (colored dots) and group-level (horizontal bars). The black error bars indicate the overall performance over conditions. The colored horizontal bar and error bar represent the mean and SEM, respectively. Val: Valence; Inf. Information. V×I: interaction between Valence and Information. ∼: .05<*P*<.1; *: .001<*P*<.01; **: .01<*P*<.001; ***: *P*<.001

Again replicating previous reports (Lebreton, et al., 2019; Salem-Garcia et al., 2021; Ting et al., 2020), our analysis confirmed that confidence, on the other hand, is additionally affected by valence (valence: F_1,39_ = 36.56, *P* < .001; information: F_1,39_ = 6.76, P = .0131; interaction: F_1,39_= 9.62, *P* =.0036). In addition to confidence being generally higher in gain than loss contexts, this valence effect was larger in the partial than in the complete information condition (post-hoc t-tests; partial: t_39_ = 6.93, *P* = 2.68×10^-8^: complete: t_39_ = 4.55, *P* = 5.08×10^-5^; difference: t_39_ = 3.10, *P* = .0451; **Figure 2B**). Overall, these results confirmed the presence of a valence-induced bias in confidence judgments that is mitigated by complete information.

We also contrasted confidence and choice accuracy to properly characterize overconfidence (or calibration). On average, calibration was non-significantly different from 0, indicating neither over-nor under-confidence (t_1,39_ = 0.1883, *P* = .8516). Yet, replicating previous finding (Lebreton, et al., 2019; Salem-Garcia et al., 2021) we found that participants were significantly overconfident in the Gain-Partial context (t_1,39_ = 2.14, *P* = .0385), and that calibration was significantly modulated by valence and information, with Losses and Complete information improving calibration (valence: F_1,39_ = 12.28, *P* = .0012; information: F_1,39_ = 14.42, *P* <.001; interaction: F_1,39_ = 0.58, *P* =.4506; **Figure 2C** and **Supplementary Table S2**). These results held when we tested generalized linear mixed-effect (GLME) models, in which we used trial-by-trial data and included predictors accounting for valence, information, the session number, and response times (**Table S3**).

Finally, response-times featured a small but significant residual effect of valence (valence: F_1,39_ = 4.77, *P* = .0350; information: F_1,39_ = 0.31, *P* = .5782; interaction: F_1,39_ = 0.97, *P* =.3318), as well as a negative correlation with confidence judgments (**Supplementary Table S2**). Despite the dissociation between decision and response processes, there was a significant correlation between response times and confidence judgments (**Supplementary Table S4**). Nevertheless, the valence-induced confidence bias and the valence-induced RT effect were not correlated at the inter-individual level (robust regression slope: β = −0.01± 0.01, *P* = .339). Moreover, an interindividual regression analysis suggested the valence-induced confidence bias could be observed in the absence of a valence-induced RT bias (robust regression intercept: β = 5.02 ± 0.84; *P* <.001; **Supplementary Table S5**). These results are in line with our previous finding that the valence-induced bias on confidence and on RTs are partially dissociable (Ting et al., 2020).

## Confidence is encoded in a positive ventromedial-prefrontal and a negative parieto-frontal network

Our neuroimaging investigations focus on confidence signals that are elicited at the decision stage (i.e., during symbol presentation, in which a motor response is not required). First, we aimed to identify neural networks whose activity generally correlates with confidence judgments during option evaluation across learning contexts. To do so, we designed a first general linear model (GLM1), in which the cue presentation period was modelled separately in each of the four contexts, and each of these events was modulated by the time series of context-specific, trial-by-trial confidence judgments (see **Methods** and **Table 1** for the complete GLM1 specification). A random-effects analysis looking at BOLD signals that were correlated with the confidence parametric modulators across contexts identified two main brain networks (voxel-wise *P*_uncorrected_ < .001; cluster-wise *P*_FWE_ < .05; **Figure 3A and Supplementary Table S6-S7**). On the one hand, neural activity in the VMPFC, pgACC, precentral gyrus, and middle temporal gyrus correlated positively with confidence rating. On the other hand, activity in large parieto-frontal network encompassing dorsolateral (bilateral IFG and INS) and dorsomedial prefrontal clusters (dACC and DMPFC) correlated negatively with confidence judgments. A small cluster in the left caudate also correlated negatively with confidence (see **Supplementary Table S7**). At the whole brain level, no brain region exhibited a valence or information effect on confidence encoding, nor an interaction between those factors (rmANOVA and direct contrasts).

**Figure 3.**
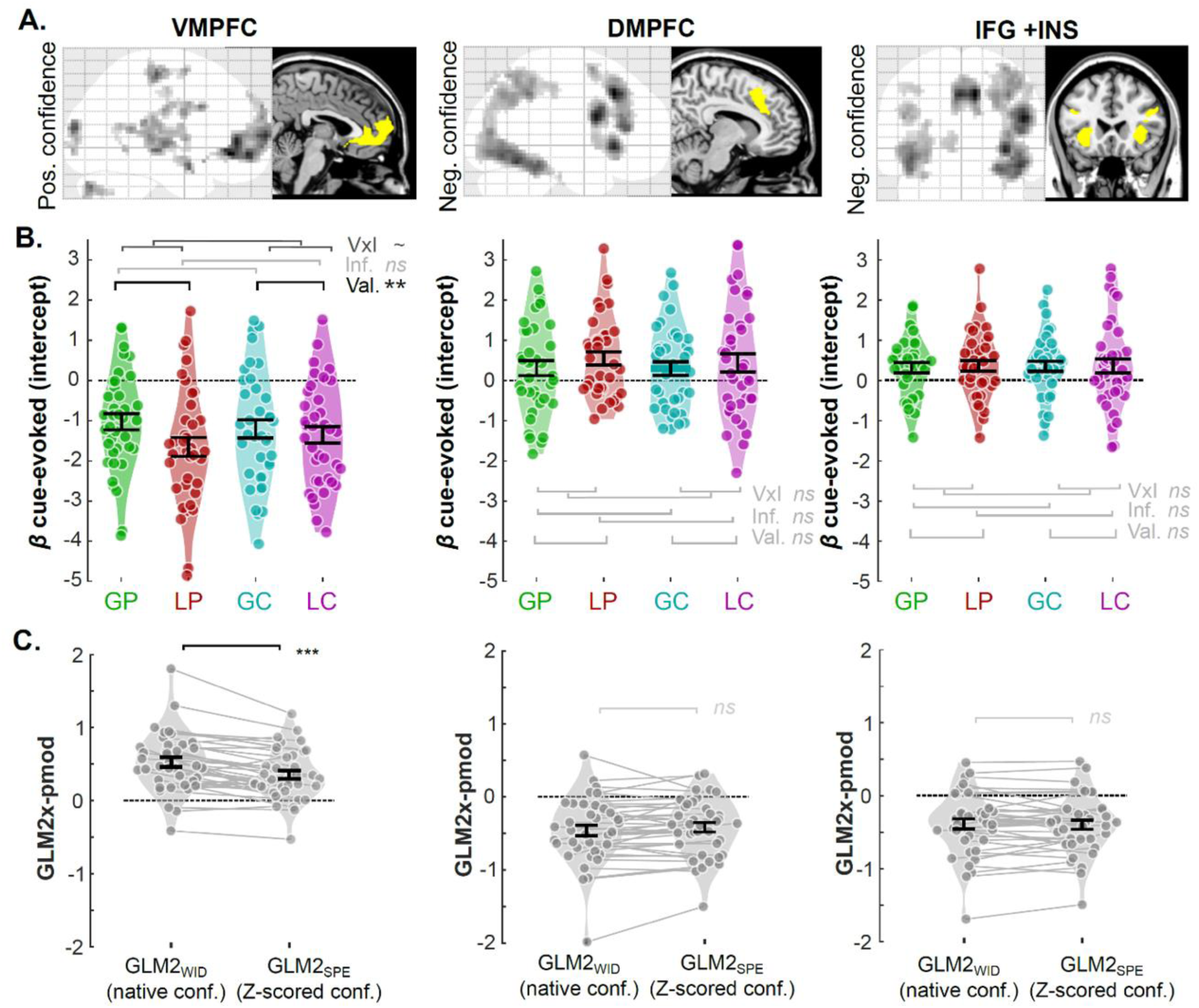
Model-free fMRI results for the learning task. **(A)** Results of whole-brain analysis. Brain areas positively (left panels) and negatively (middle and right panels) correlate with confidence rating during the symbol presentation phase. Significant voxels are displayed on the glass brains in a gray-to-black gradient manner (*p*uncorrect< .001, cluster size >47). The yellow areas in the anatomical brain are ROIs (vmPFC, dmPFC, and IFG+INS), which are used in the following ROI analyses. **(B)** Violin plots represent the sample distribution of fMRI regression coefficients of cue-evoked signals for the different contexts (represented by different colors). Dots correspond to individual regression coefficients. Error bars represent sample mean ± SEM. GP: gain/partial; LP: loss/partial; GC: gain/complete; LC: loss/complete. **(C)** Violin plots represent the sample distribution of fMRI regression coefficients for native versus Z-scored confidence regression coefficients, respectively extracted from GM2WID and GLM2SPE. Dots correspond to individual regression coefficients. ∼: .05<*P*<.1; *: .001<*P*<.01; **: .01<*P*<.001; ***: *P*<.001

**Table 1.**
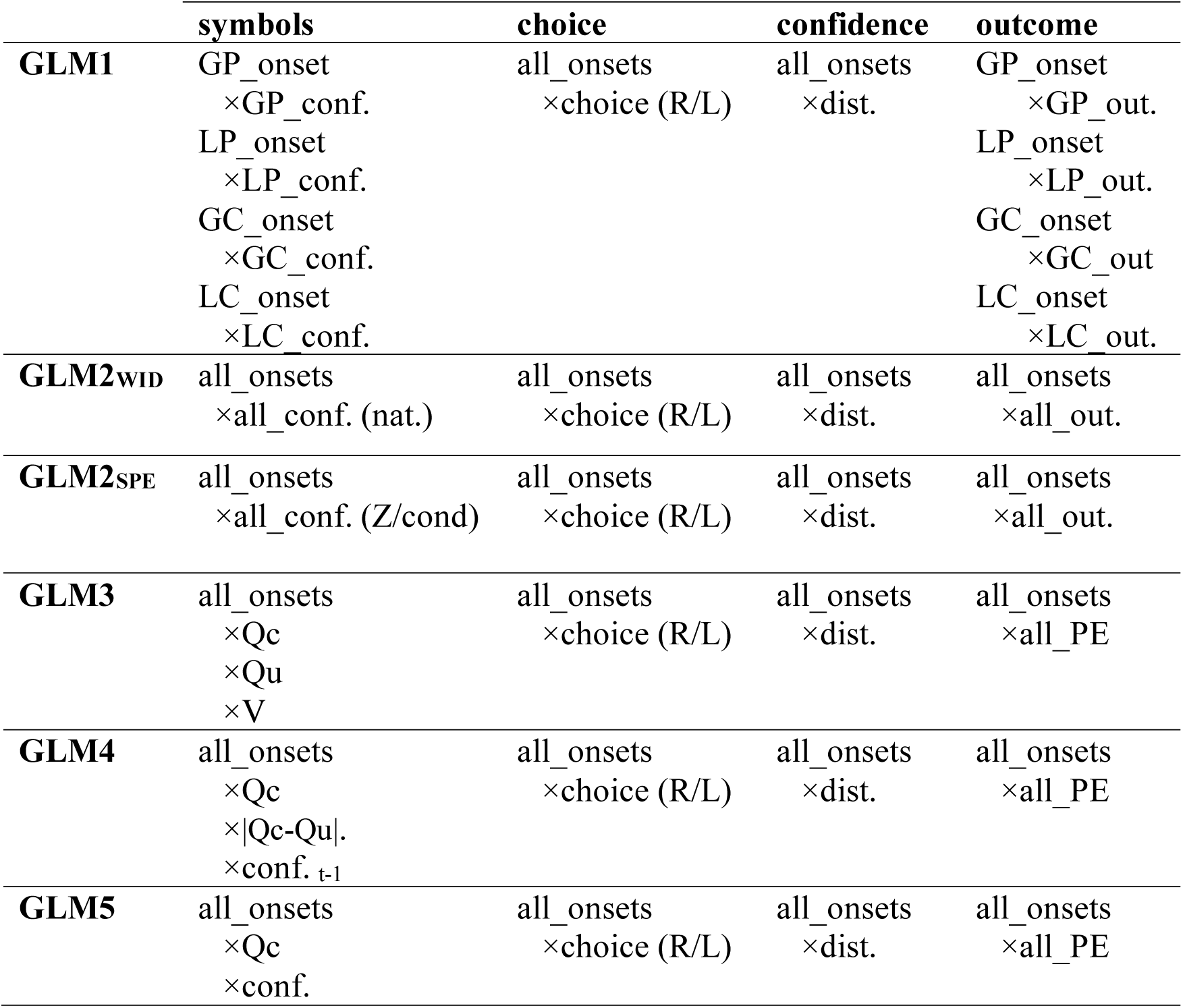
GLMs’ structure. The table represents the four events of interest in a trial as columns, and list for each GLM, the corresponding regressors and their respective parametric modulators (indicated by a × sign). GP: Gain-Partial; LP: Loss-Partial; GC: Gain-Complete; LC: Loss-Complete conf: confidence; (R/L): choice coded as 1/-1 for right/left. dist.: distance (difference between the starting point and final confidence rating); out.: outcome (coded 1/0 if the chosen outcome is the best/worst potential outcome ‒ i.e., 1 and −0.1 are encoded as 1 and 0.1 and −1 are encoded as −1). Qc: chosen option values. Qu: unchosen option value. V: context value. PE: prediction error. Parametric modulators, Qc, Qu, V, and PE, are estimated by the winning model.

To better characterize the signal encoded in the confidence-encoding prefrontal regions, we then regrouped the prefrontal clusters identified in our whole-brain analysis into three main functional regions(/networks)-of interest (ROIs), respectively representative of ventromedial (VMPFC), dorsolateral (DLPFC: union of bilateral INS and IFG) and dorsomedial (DMPFC, dACC) prefrontal cortices. Then, we extracted, in these ROIs, the parametric confidence regression coefficients for all four contexts. We first verified that our experimental manipulations of outcome valence and outcome information did not impact this parametric encoding of confidence (all Ps > 0.05/3; **Figure S6B** and **Supplementary Table S6-S7**). No significant effect of those factors was found (Bonferroni-corrected for three comparisons). Overall, these analyses confirmed that VMPFC on the one hand, and DLPFC and DMPFC one the other hand, respectively constitute the positive and negative confidence-encoding networks, and that they encode confidence similarly across the different context.

## Task-wide vs. condition-specific confidence in the brain

Next, we turned to our main question of interest, namely dissociating different types of confidence and uncertainty signals, which we ultimately hoped could help in identifying functionally dissociable brain networks. We defined three theoretical types of qualitative patterns on those cue-evoked activities, that specifically characterize three confidence related neural signals: uncertainty, condition-specific confidence and task-wide confidence (**Figure 1D**). Essentially, statistical uncertainty corresponds to the objective difficulty of the choice, that is ultimately revealed in choice accuracy. Accordingly, statistical uncertainty should be higher in Partial than in Complete information contexts, but identical in Gain and Loss contexts, given the similar objective difficulty and observed performance between these conditions (**Figure 1D**). Condition-specific confidence simply tracks the subjective, relative improvement within each context, and is reminiscent of the context value that tracks the choice-independent expected value in each context (Palminteri et al., 2015; Palminteri & Lebreton, 2021). Thereby, condition-specific confidence should be purely context dependent, hence not show any effect of our factors (**Figure 1D**). Finally, task-wide confidence corresponds to the actual absolute, phenomenological feeling of confidence that is reported as the confidence judgments. Task-wide confidence should then be higher in Gain than Loss context, with potentially a mitigation by information (**Figure 1D**). From those definitions, and given that our ROIs have already been shown to encode confidence across contexts, one can simply ascribe those theoretical variables to ROI activity, by testing the effect of valence and information on cue-evoked activity, as modelled in GLM1 (**Figure 1D**). We found a significant valence effect (F_1,37_ = 8.99, *P* = .0048) and marginal valence-information interaction in VMPFC (F_1,37_ = 4.22, *P* = .0532) (**Figure 3B and Supplementary Table S6**). Mimicking the pattern of confidence judgments, the difference between BOLD activity elicited in gain versus loss contexts was higher in the partial than in the complete information context (partial: 0.62±0.18; complete: 0.14±0.16; t_37_ = 1.99, *P* = .0532). In contrast, we did not find significant effects of the valence and information factors on BOLD activity in either of the negative networks (*Ps* > 0.8). The results of this ROI analysis tentatively ascribe task-wide confidence signals (including a valence effect) to the VMPFC and condition-specific confidence (without valence nor information effects) to both DLPFC and DMPFC. For completeness, we also tested for additional whole-brain activation for the positive and negative effects of valence and information on cue-evoked activity. The result revealed that only the Gain > Loss contrast elicited activations in a large brain network encompassing, among other regions, the VMPFC (voxel-wise *P*_uncorrected_ < .001; cluster-wise *P*_FWE_ < .05; **Figure S6** and **Supplementary Table S7**). Finally, we performed a whole-brain conjunction between regions correlating positively with confidence and regions positively encoding valence (i.e., Gain > Loss). Again, we found that BOLD signal in the VMPFC jointly correlated with valence and confidence, suggesting that it plays a key role in processing a global, task-wide confidence signal (voxel-wise *P*_uncorrected_ < .001; cluster-wise *P*_FWE_ < .05; **Figure S6C**).

## Quantitative assessment of confidence-related variable encoding

Although the analysis of the qualitative patterns of activations seem to clearly point to a functional dissociation between the positive and negative prefrontal network in confidence encoding, some aspects of the demonstration still have some weaknesses. For instance, ascribing condition-specific rather than task-wide confidence signal to the negative network entails accepting the null hypothesis – i.e. concluding that valence and information are not statistically detectable in the negative network ROIs’ signal. Here, we propose a different set of analyses to quantitively support this conclusion without relying on this statistical caveat. To provide a fair comparison between task-wide and condition-specific confidence, we designed two new GLMs (GLM2_WID_ and GLM2_SPE_), that concatenated all learning-contexts into one single cue-evoked event (i.e., symbol presentation period). Then, in GLM2_WID_, this event was modulated by the time series of all *native* confidence judgments (i.e., the absolute confidence reports provided by our subjects on each trial). On the contrary, in GLM2_SPE_, this event was modulated by the time-series of all confidence judgments, but normalized (i.e. Z-scored) per condition (i.e., reflecting variation around each condition mean). This way, the structure of these two GLMs is identical, but the parametric modulators of confidence respectively represent task-wide confidence (i.e., native, absolute confidence) and condition-specific confidence. We then extracted the confidence regression coefficients from our ROIs, and proceeded to two types of quantitative comparisons. First, we simply compared the GLM2_SPE_ and GLM2_WID_ regression coefficients (**Figure 3C**). In the VMPFC, activations related to native confidence were significantly higher than to normalized confidence (t_37_ = 5.41, *P* <.001). In total, this pattern was found in 30 out of 38 participants, further evidencing that activity in the VMPFC better corresponds to task-wide than condition-specific confidence. However, the same analysis was inconclusive for the regions of the negative network –although trending in the direction of higher activations for condition-specific confidence for some regions (DMPFC: t_37_ = −1.40, P = .1670; IFG+INS: t_37_ = 0.43, P = .6684). Note that the underlying test that was used to create ROIs, a grouping parametric effect of confidence from GLM1, was orthogonal to the follow-up tests on task-wide and condition-specific confidence encoding, therefore these analyses were not circular and did not advantage GLM2_SPE_ or GLM2_WID_ (Kriegeskorte et al., 2009). We then complemented analysis with a formal Bayesian Model Comparison (BMC; see **Supplementary Methods: Bayesian model comparison**) between the GLM2_SPE_ and GLM2_WID_ in our ROIs, using the SPM-based MACS toolbox (Soch & Allefeld, 2018). This time, while the analysis was inconclusive in the VMPFC (GLM2_WID_ vs GLM2_SPE_; Exceedance Probability EP: 48.69% vs 51.31%), it provided suggestive evidence that lateral and dorsal parts of the negative network are better explained by condition-specific than task-wide confidence (GLM2_WID_ vs GLM2_SPE_ EP: DMPFC: 17.78% vs 82.22%; IFG+INS: 09.66% vs 90.34%). Overall, converging evidence from different models and statistical tools seem to confirm our functional dissociation between the VMPFC and the negative network.

## Computational models for learning and confidence judgments

The vast majority of past studies investigating neurocomputational models of reinforcement-learning have focused on the neural representation of learning latent variables such as option and action values, prediction errors, and various levels of (Bayesian) uncertainty. As a matter of fact, the emerging consensus in the RL literature seems to indicate that neural signal in the VMPFC is specifically linked to the representation of option values, from which decisions are derived (Liu et al., 2011; Rushworth et al., 2011). Evaluating the relative merits of our current hypothesis against this consensus, namely that VMPFC encodes confidence judgments rather than values during RL, requires a computational model that faithfully captures our participants’ behavior and that can produce the desired latent variables. Following the rationale of a recent study (Salem-Garcia et al., 2023), we proposed a combination of a RL model and of a confidence regression, to jointly account for behavioral choices and confidence judgments exhibited in the current experimental framework (i.e. in both the learning and transfer phases). We factorially tested several families of RL model (**Figure 4A** and **Methods**), which built on a basic Q-learning model (ABS), and modularly featured context-dependent learning (RELATIVE family) as well as confirmatory updating (ASYMMETRIC family) ‒ see also (Palminteri & Lebreton, 2021, 2022). Replicating previous findings, we found that both features were necessary to best account for our participant data, as revealed by a formal Bayesian Model Selection (BMS) analysis (**Figure 4B-C**; winning model: RELASYM; protected Exceedance probability >99%). The RL model provided latent variables (i.e. option Q-values and context-value V), from which we then built several confidence models (**Figure 5A** and **Methods**). Confidence models consisted in a logit-transformed multiple regressions that included, as predictor variables, choice difficulty –proxied by the absolute difference between option values (|Qc−Qu|)–, plus a biasing term accounting for the valence-induced bias (for which we tested several variants: 0, ΣQ, V, Qc; **Figure 5A**), and an autocorrelation term (i.e. confidence in the previous trial) that accounts for the tendency of confidence judgments to exhibit serial dependency (Rahnev et al., 2015). A BMS revealed that the confidence model that featured the value of the chosen option Qc as a biasing term (thereafter referred to as Qc-REG) provides the best account of participants confidence judgments (protected Exceedance probability >99%; **Figure 5B-C**). In the supplementary methods of the present paper (**Supplementary Figure S1-S5**), we systematically apply the set of analyses underlying the demonstration proposed in (Salem-Garcia et al., 2023) and compare its results to those obtained in the present dataset (learning + transfer). This exercise confirmed that the combination of RELASYM and Qc-REG models faithfully capture our participants’ behavior (choice and confidence judgments) throughout our experimental framework (learning and transfer phase), and that learning biases are fundamentally linked with confidence biases (**Supplementary Figure S5**).

**Figure 4.**
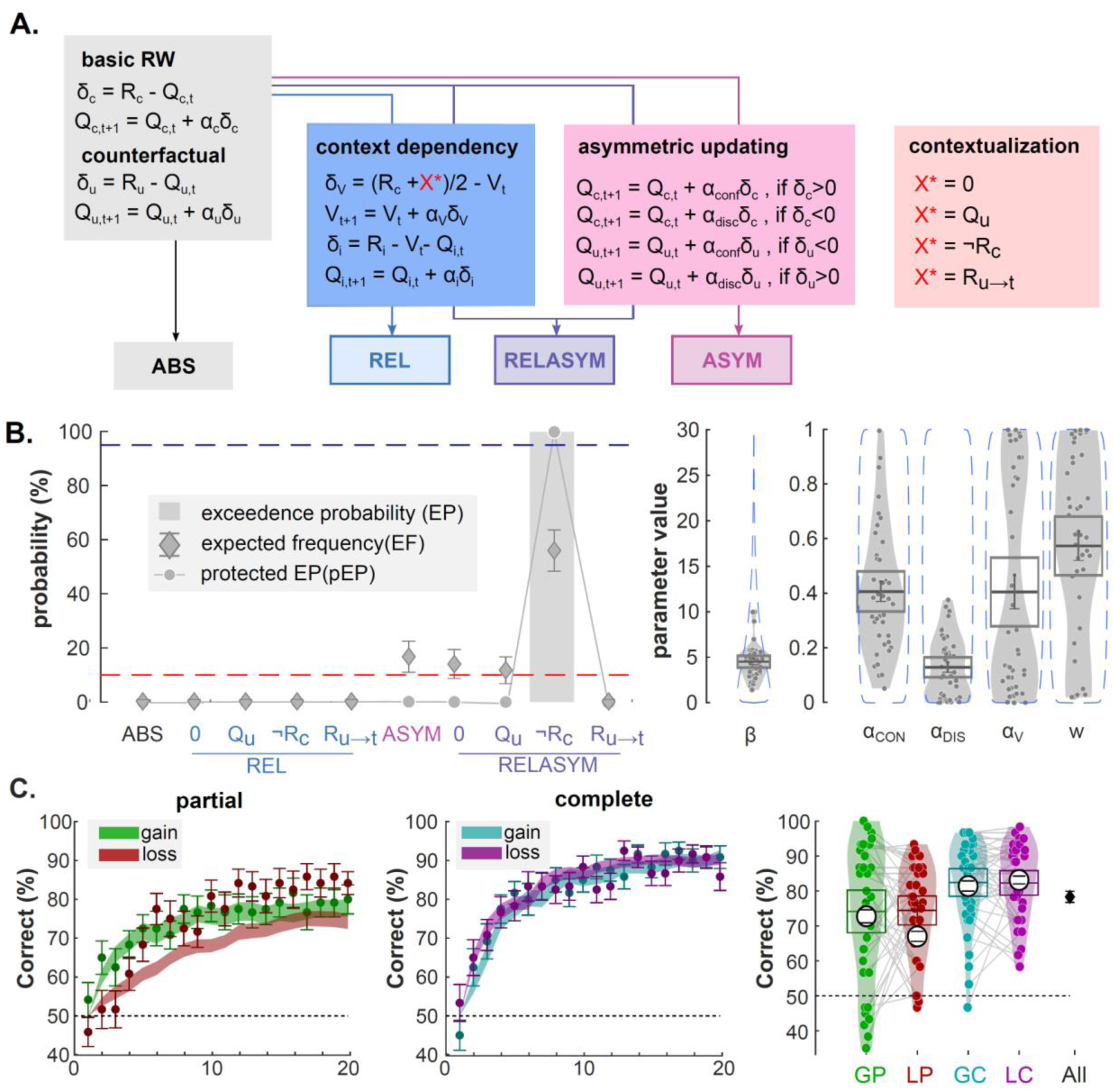
Modeling choices in the learning phase. **(A)** The learning model architecture to explain participants choice data. Color panels represent different components of value updating rules. Gray panel: Absolute model (ABS), which consists of basic Rescorla-Wagner (RW) update rule. This rule updates chosen and unchosen option values via outcome directly. Blue panel: Relative model (REL), which consists context-dependent component and updates option values by considering context value. Pink panel: Asymmetric updating model (ASYM), which updates option values based on the valence of prediction error. Purple panel: relative-asymmetric model (RELASYM), which is the combination of relative model and asymmetric updating model. The contextualization panel is used to update unchosen option in the partial information condition. Specifically, X* is determined as the unchosen option outcome (Ru) when the value is available in the complete information condition. When the unchosen option outcome (Ru) is not available in the partial information condition, X* is hypothesized as none (0), expected unchosen value (Qu), paired outcome (¬R) and last seen outcome associated with the option (Ru→t). **(B)** Left panels: Bayesian model comparison. X-axis represents the models with different hypothesized contextualization values. Y-axis represents the value of three criteria, including exceedance probability (EP; grey histograms), expected frequencies (EF; diamonds) and protected exceedance probability (pEP; line & dots) of each model. The red dashed line represents the guessing level for EF. The blue dashed line represents the threshold (95%) for the exceedance probability. Right panels: Estimated parameter values of the winning model (RELASYM, X* = with ¬Rc). Dots represent individual data points. Error bars displayed within the violin plots indicate the sample mean ± SEM. The blue, dotted envelop represent the prior distribution. **(C)** Left: modelled trial-by-trial percentage of correct responses. Dots and error bars represent the mean ± SEM of the participant data. Filled, shaded colored areas represent mean ± SEM of the posterior predictive fits obtained from our winning computational model (RELASYM, X* = with ¬Rc). Right: average percentage of correct responses across conditions at the individual level (colored dots) and group-level (horizontal bars). The black error bars indicate the overall performance over conditions. The colored horizontal bar and error bar represent the mean and SEM, respectively. The large with dot and corresponding error bar represent mean±SEM of the posterior predictive fits obtained from our winning computational model (RELASYM, X* = with ¬Rc). Qc/u,t: value of the chosen/unchosen option at trial t. Rc/u: outcome associated to the chosen/unchosen option. δc/u: prediction error for the chosen/unchosen option. αu/c: learning rate for the chosen/unchosen option. αconf/disc: learning rate for confirmatory/disconfirmatory information. Vt: context value; δV: prediction error for the context value. αV: learning rate for the context value.

**Figure 5.**
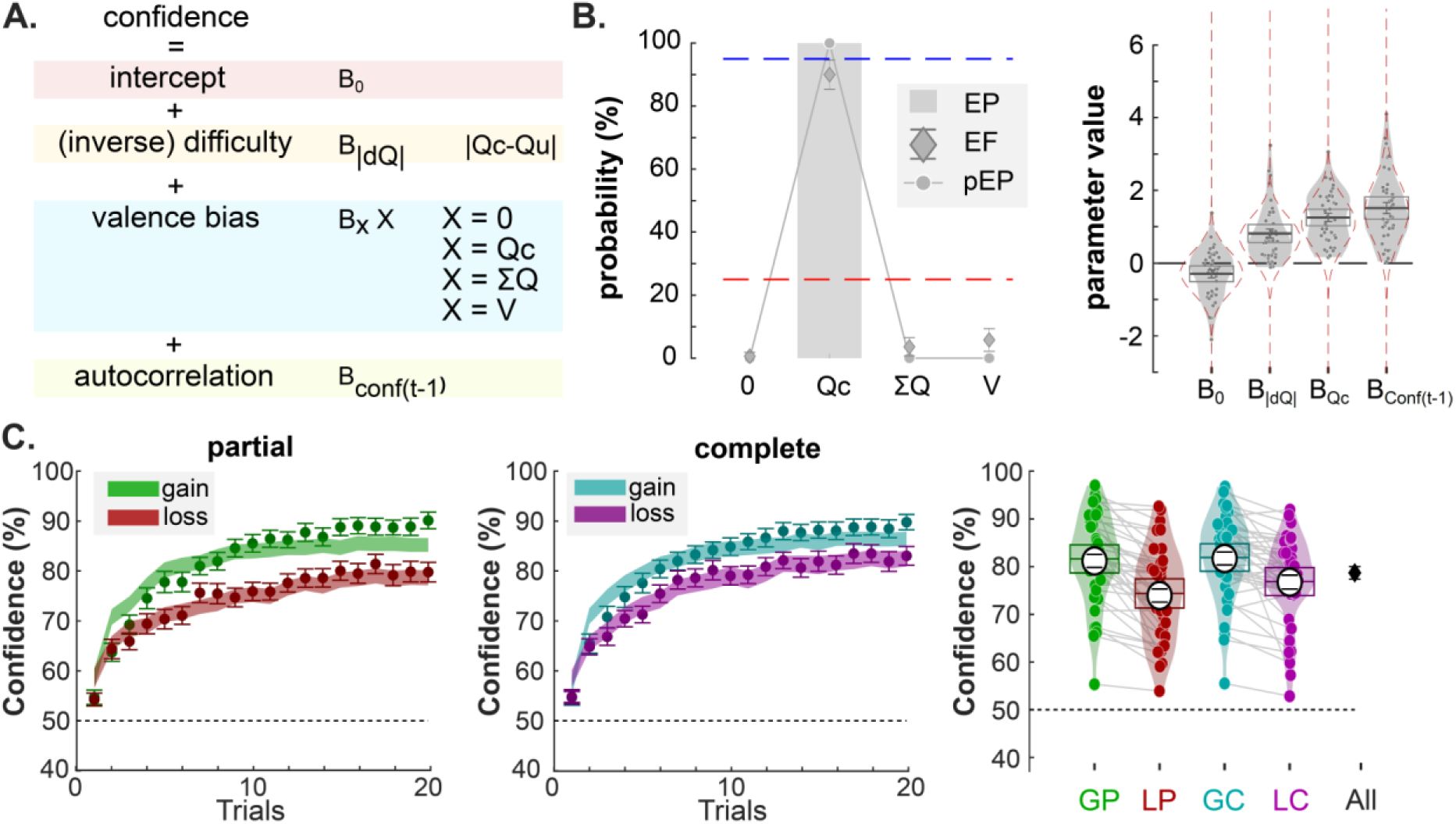
Modeling confidence in the learning phase. **(A)** The confidence model architecture to explain participants confidence judgment data. Color panels represent different components of the multiple regression predicting confidence. In particular, the blue rectangle pictures the different hypotheses for the biasing term. **(B)** Left panels: Bayesian model comparison. X-axis represents the models with different hypothesized valence biases. Y-axis represents the value of three criteria, including exceedance probability (EP; grey histograms), expected frequencies (EF; diamonds) and protected exceedance probability (pEP; line & dots) of each model. The red dashed line represents the guessing level for EF. The blue dashed line represents the threshold (95%) for the exceedance probability. Right panels: Estimated parameter values of the winning model (Qc-REG). Dots represent individual data points. Error bars displayed within the violin plots indicate the sample mean ± SEM. The blue, dotted envelop represent the prior distribution. **(C)** Left: modelled trial-by-trial confidence judgments. Dots and error bars represent the mean ± SEM of the participant data. Filled, shaded colored areas represent mean ± SEM of the posterior predictive fits obtained from our winning model (Qc-REG). Right: average confidence across conditions at the individual level (colored dots) and group-level (horizontal bars). The black error bars indicate the overall performance average across conditions. The colored horizontal bar and error bar represent the mean and SEM, respectively. . The large white dots and corresponding error bar represent mean ± SEM of the posterior predictive fits obtained from our winning computational model (Qc-REG). Qc/u: value of the chosen/unchosen option; V: context value; ΣQ: sum of chosen and unchosen Q-values

## BOLD activity in the positive and negative networks correlate with decision values

Thanks to the latent variables estimated form our computational models, we next tested whether activity in the prefrontal regions originally identified in our confidence analyses (**Figure 3A**; VMPFC; DMPFC; IFG+INS) could also be explained with the more traditional learning and decision variables. We therefore designed a new GLM (i.e., GLM3, see **Table 1**) for a model-based fMRI analysis, which comprised, as parametric regressors of the cue onset, all value-related latent variables estimated by the RELASYM model: the chosen option value (Qc), the unchosen option value (Qu), and the context value (V). We then extracted the parametric regressors in the three main regions forming our confidence networks. Altogether, and in line with previous findings (Liu et al., 2011; Palminteri et al., 2015; Rushworth et al., 2011), we found that the chosen option values (Qc) correlated with BOLD activity positively in the VMPFC (t_37_ = 3.26, *P* =.0023) and negatively in the DMPFC (t_37_ = −4.96, *P* < .001) and IFG+INS (t_37_ = −4.43, *P* < .001) (**Figure 6A**). In addition, the unchosen option value (Qu), correlated positively with BOLD activity in the DMPFC (t_37_ = 2.96, *P* = .0053) and IFG+INS (t_37_ = 2.75, *P* = .0091). At the whole-brain level (*P*_FWE_ < 0.05 at the cluster level), only the chosen option values (Qc) generated significant clusters of activations in the prefrontal regions, in both the VMPFC (positive) and in the IFG+INS (**Supplementary Table S8).** Therefore, in the context of reinforcement learning, neural activity in the ventral and dorsal prefrontal cortices can be evenly ascribed to two very different cognitive processes: the computation of decision values and/or the evaluation of confidence in the upcoming decision.

**Figure 6.**
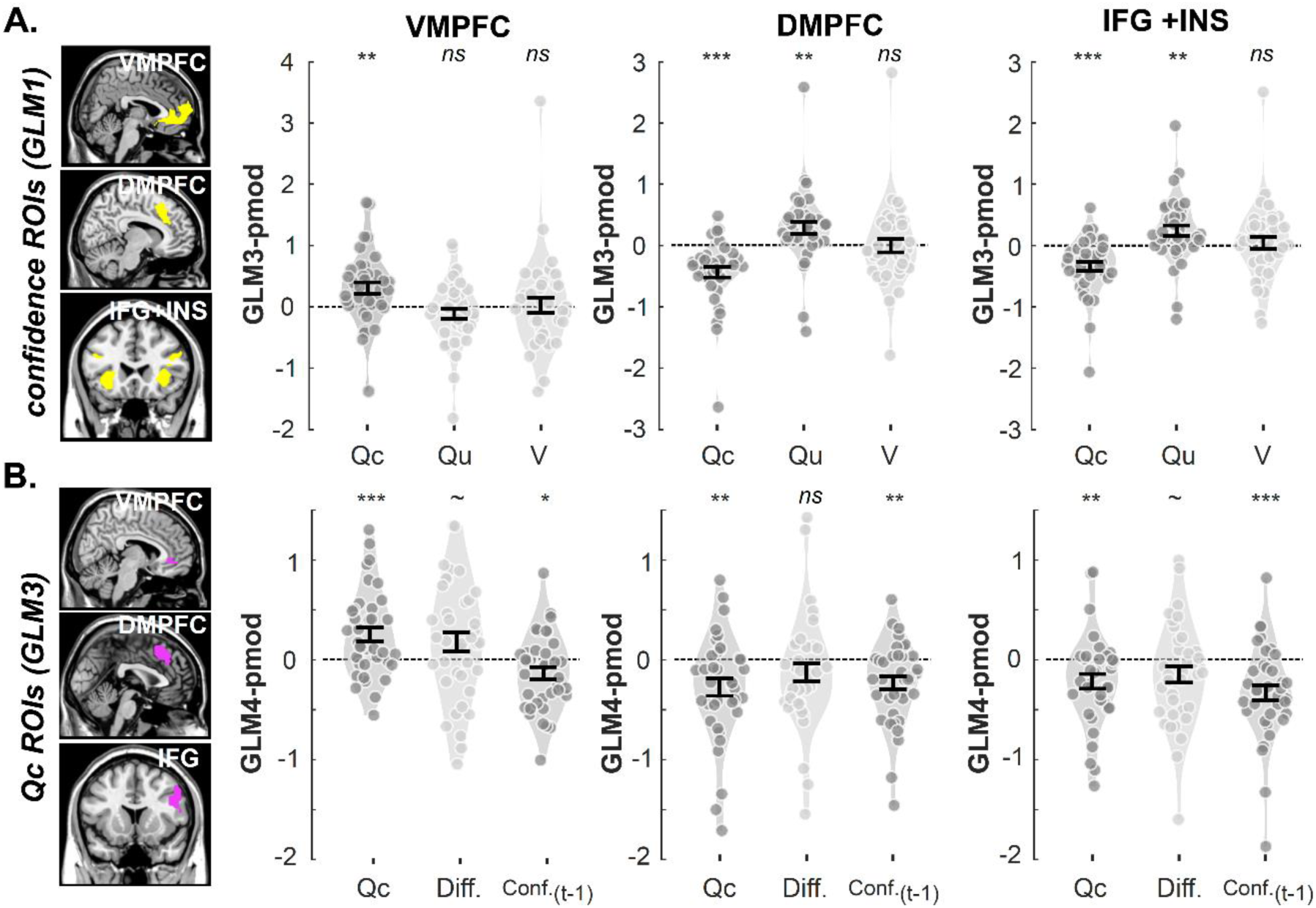
vmPFC is involved in value and confidence processing. Violin plots represent the sample distribution of fMRI regression coefficients corresponding to several variable of interest included in different GLMs, extracted from each ROI (left: VMPFC; middle: DMPFC; right: IFG+INS) at the symbol presentation phase. (A) Regression coefficients for RL-derived value latent variable. Dots correspond to individual regression coefficients. (B) Regression coefficients for confidence model latent variables. ROIs were defined with confidence activations from GLM1 (A) or Qc-activations from GLM3 (B). Dots correspond to individual regression coefficients. Dark gray and light gray indicate the effect is significantly and insignificantly different from 0, respectively. Error bars represent sample mean±SEM. **Qc**: parametric modulator of chosen option; **Qu**: parametric modulator of chosen option.; **V**: parametric modulator of context value.; **Diff.**: parametric modulator of absolute value difference of Qc and Qu. ∼: .05<*p*<.1; *: .001<*p*<.01; **: .01<*p*<.001; ***: *p*<.001

## BOLD signal in the VMPFC correlates with confidence-building variables

To evaluate whether prefrontal activations with confidence could have been purely confounded (i.e., explained) by their role in computing decision values (notably Qc, in the VMPFC), we proceeded to a reverse double dipping exercise. We created a new GLM (i.e., GLM4), which contained the three components of confidence suggested by the Qc-REG models (Qc, |Qc−Qu|, and conf._t-1_) as parametric regressors of the cue onset. We defined as our three prefrontal ROIs the significant clusters revealed by the whole brain correlation with Qc in GLM3, and extracted the parametric regressors of the confidence component estimated with GLM4. Critically, the VMPFC ROI that was selected to be specifically associated with Qc also exhibited residual correlations with the other confidence components (**Figure 6B;** Qc: t_37_ = 3.63, *P* < .001; |Qc−Qu|: t_37_ = −3.19, *P* = .0029; conf._t-1_: t_37_ = −2.93, *P* = .0057). Note, however, that when the different sources of confidence formation competed for the variance of BOLD signals, only Qc elicits whole-brain significant activations in VMPFC (voxel-wise *P*_uncorrected_<.001; cluster-wise *P*_FWE_ <.05; **Supplementary Table S9)**. Still, those analyses suggest that the VMPFC does not simply encode Qc, but exhibit additional signatures of confidence signal.

## BOLD signal in the VMPFC is better explained by confidence than decision variables

A recent stream of studies has suggested that, in simple decision-making or judgment situations, the VMPFC encodes a combination of both decision values and confidence (De Martino et al., 2013; De Martino et al., 2017; Lebreton et al., 2015a; Lopez-Persem et al., 2020). In this last section, to refine the characterization of VMPFC activity during human reinforcement-learning, we estimated an fMRI model which included both Qc and confidence judgments as parametric regressors (GLM5). Following the rationale of (Lebreton et al., 2015), we designed value-related VMPFC ROIs, from the Qc-activations revealed in GLM3, and from a meta-analysis of fMRI activations value (Bartra et al., 2013). We then extracted regression coefficients of Qc and confidence from the GLM5 model, so as to test for the presence of confidence signals in those value-coding regions (**Figure 7A**). Despite the choice of our ROIs, which should bias our analyses in favor of value activations, the Qc-related activations were marginal to insignificant (**Figure 7A**, GLM3-ROI: *P* = .0553; Bartra ROI: *P* = .2324) in our model in which value- and confidence-related parametric modulators compete for variance. On the contrary, confidence-related activations were clearly significant in both ROIs (**Figure 7A**, *Ps*<.0001), and significantly larger than Qc-related activations (**Figure 7A**, *Ps*<.05). Note that a formal comparison between models featuring one (Qc or confidence) versus two (Qc and confidence) using BMC failed to provide conclusive results. In the negative network (DMPFC; INS+IFG), the comparison of confidence and Qc-parametric regressors did not reach significance, again suggesting a functional dissociation with its positive counterpart (**Supplementary Figure S7**).

**Figure 7.**
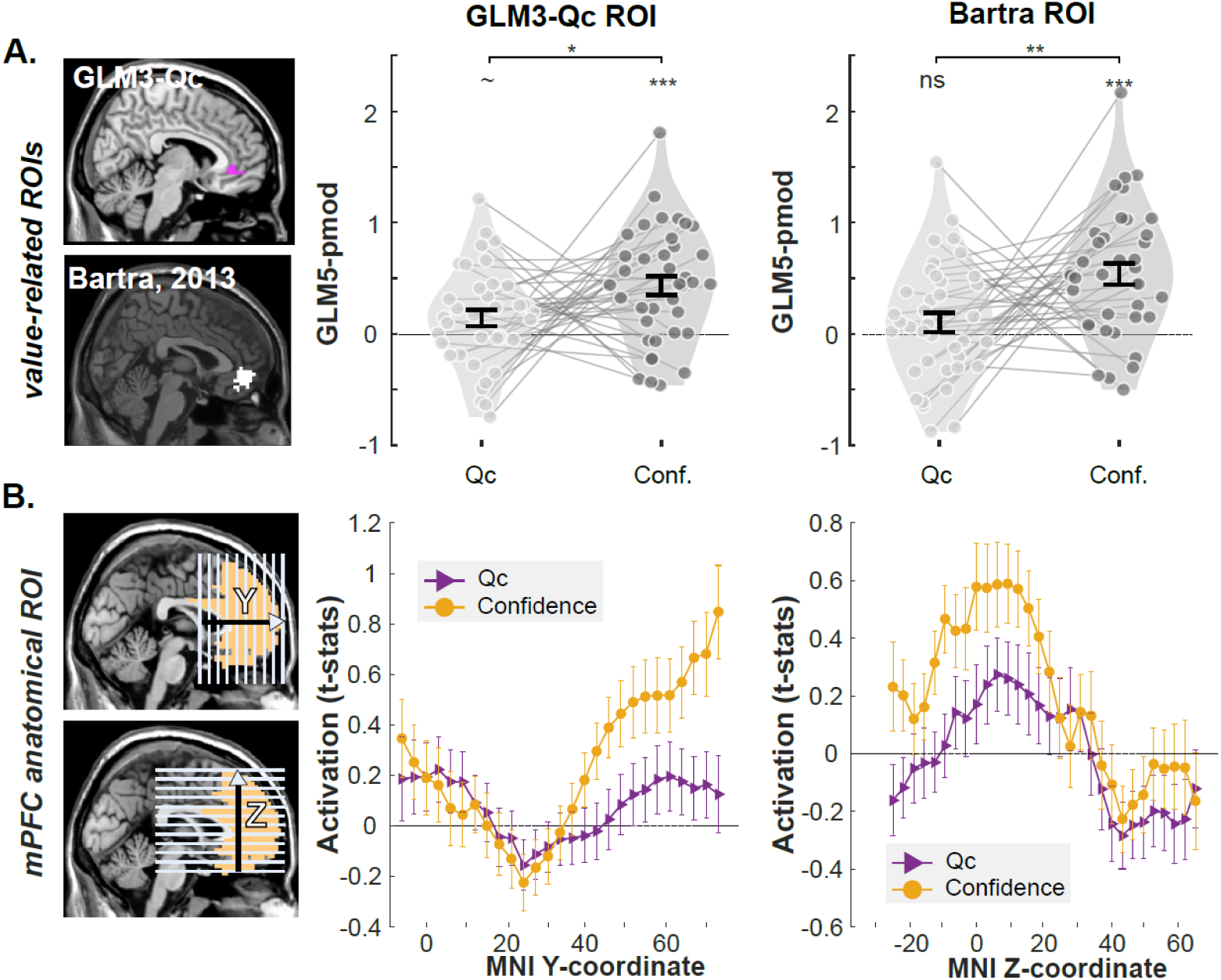
value and confidence activations in the VMPFC. **(A)** ROI analysis with Qc-related ROIs identified in the present study (top-left; purple areas) and in an independent meta-analysis (bottom_left; white area). Right: the regression coefficients corresponding to Qc and confidence in GLM5 were summarized at the individual level (dots). Violin plots represent the sample distribution of fMRI regression coefficients. Dots correspond to individual regression coefficients **(B)** Anatomical ROI of mFPC. BOLD signal was extracted along y-dimension from posterior to anterior area and along z-dimension from ventral to dorsal area (pictured slices are only illustrative, and do not indicate the actual coordinate of the extracted signal). Voxel-wise t-values of Qc and confidence in GLM5 were extracted and averaged over two dimensions. Middle : average t-value along MNI y-coordinate. Right: average t-value along MNI y-coordinate. Dots and error bars represent mean ±SEM. **Qc**: parametric modulator of chosen option; **Conf.**: parametric modulator of confidence ratings ∼: .05<*P*<.1; *: .001<*P*<.01; **: .01<*P*<.001; ***: *P*<.001

Finally, we considered the possibility that value and confidence signals dominate in different sub-regions of the prefrontal cortex (Clairis & Pessiglione, 2022). Therefore, following the rationale in (Clairis & Pessiglione, 2022; Hoven, et al., 2022), instead of averaging signal over the entire ROI, we extracted regression coefficients in a large anatomical prefrontal ROI, and marginalized those activations along the anterio-posterior (Y) and ventro-dorsal (Z) axes (**Figure 7B**). This finer-grained analysis revealed that confidence activations dominate value-activations over all portions of the medial prefrontal cortex.

## Discussion

Decisions are usually accompanied by confidence judgements, which reflect subjective (un)certainty about the choice being correct (Fleming & Dolan, 2012; Lebreton et al., 2015; Pouget et al., 2016). This internal signal plays a crucial role in guiding behaviors and has been associated with two main prefrontal networks: VMPFC and DMPFC (Bang & Fleming, 2018; Rouault, Lebreton, et al., 2022; Vaccaro & Fleming, 2018). To date, though, the relative contribution of those two networks in the mechanisms underlying confidence formation remains unclear. To fill this gap, we combined fMRI and an adapted probabilistic reinforcement learning task (Lebreton, et al., 2019; Palminteri et al., 2015; Ting et al., 2020), in which we systematically manipulated two dimensions of the learning context: the valence of the outcome (gain vs. loss) and the outcome information (partial vs. complete feedback). At the behavioral level, we successfully replicated the valence effect on confidence judgments: confidence is significantly higher when learning to gain rewards relative to learning to avoid losses, despite participants learning equally well in both contexts (Lebreton, et al., 2019; Salem-Garcia et al., 2021; Ting et al., 2020). At the neural level, we first replicated consensual and established results: confidence was positively related to the activation in the VMPFC and neighboring area pgACC (positive-confidence network) and negatively related to the activation in the DMPFC, IFG, and INS (negative-confidence network) (Bang & Fleming, 2018; Rouault, Lebreton, et al., 2022; Vaccaro & Fleming, 2018). Then, we uncovered two new key findings. First, our analyses revealed that VMPFC activity represents a task-wide (subjective) confidence signal as it tracks confidence within contexts together with the valence bias that increases confidence in gain contexts. Activation in the negative-confidence network (DLPFC, DMPFC), on the other hand, only tracks condition-specific confidence. Accordingly, we speculated that the VMPFC is a key region involved in the valence-induced confidence bias during reinforcement learning. Second, we found that, contrary to the current dominant view in the field, the activation in the VMPFC can be better explained by confidence rather than other value-related variables estimated by a RL model. In the following sections, we discuss these new findings in more details.

The simultaneous neural representation of valence and confidence in the VMPFC suggests that VMPFC integrates affective/motivational information with metacognition, and as such plays a key role in the valence-induced confidence bias (De Martino et al., 2013; De Martino et al., 2017; Fleming et al., 2012, 2014; Hoven, et al., 2022; Hoven, et al., 2022; Lebreton et al., 2015, 2018). Contrary to our theoretical predictions, we did not identify a brain region that is sensitive to confidence and to the information manipulation (i.e., partial and complete feedback). This might be due to the low effect size of information on confidence (though effects on accuracy are clear) or the fact that, as our modelling suggests, participants tend to infer the counterfactual outcome when not observed ‒ see **Figure 4** and (Salem-Garcia et al., 2023; Ting et al., 2021). Another possibility is that, while confidence-related variables are explicitly monitored by some brain areas, uncertainty is implicitly encoded in the variance of neural populations, which our current neuroimaging approach would fail to capture (Knill & Pouget, 2004; van Bergen et al., 2015).

In addition, our results provide evidence for the co-existence of task-wide confidence in VMPFC and condition-specific confidence in DMPFC. This functional difference confirms that those two brain networks are not redundant in the way they process confidence-relevant information (Bang & Fleming, 2018; Rouault, Lebreton, et al., 2022), but also raise legitimate questions about the advantages of tracking both variables and the relation between them. Naturally, access to task-wide (i.e. absolute) confidence is critical to compare (or even choose between) different choice situations whose assessment regarding the probability of being correct differ (de Gardelle & Mamassian, 2014). Task-wide confidence can be viewed as an overarching estimate of confidence that enables to select situations in which we perform well, and avoid situations in which we perform less well. Its role of monitoring confidence across multiple contexts therefore places task-wide confidence in an advantageous position to solve the explore exploit problem. Yet, evidence suggest that most neural and cognitive computations are context-dependent (Carandini & Heeger, 2012; Louie & De Martino, 2014), notably in the context of reinforcement learning (Hunter & Daw, 2021; Palminteri & Lebreton, 2021), such that metacognition and confidence might not elude this general neurocomputational principle. While our current results remain agnostic about the mechanistic interactions between task-wide and condition-specific confidence, most models of confidence formation seem to assume that local variables (e.g. uncertainty or condition-specific confidence) are precursors of more general, absolute confidence judgments (Cortese, 2021; Pouget et al., 2016; Rouault et al., 2019). In our case, this would imply that early, condition-specific signals in the negative network (DMPFC, DLPFC) are then fed to the positive network (VMPFC), where a general, task-wide confidence signal matches the report of participants which corresponds to the subjective experience -i.e. phenomenological dimension- of the feeling of confidence (Bang & Fleming, 2018; Lau et al., 2022) ‒ but see (Gherman & Philiastides, 2018) for evidence of opposite patterns. Finally, a couple of recent studies investigated how a global feeling of confidence (over a whole task) builds from multiple local signals (over trial-by-trial changes in task difficulty and performance), and report that VMPFC tracks local confidence in a manner that is sensitive to global self-performance estimations (Rouault et al., 2019; Rouault & Fleming, 2020). Altogether these results seem to indicate that VMPFC aggregates a complex confidence estimates over multiple layers of precursor variables.

Two main lines of arguments motivated us to complement our first set of neuroimaging analyses focused on confidence signals with model-based assessments of value-related signals. First, similarly to the decision-making literature, the reinforcement-learning literature has so far mostly associated VMPFC with the processing of value – rather than confidence (Liu et al., 2011; Rushworth et al., 2011). Second, we recently suggested that, during reinforcement-learning, confidence builds notably on two variables estimated from learned option-values: the choice difficulty (proxied by the absolute difference between the two available options values), and the chosen option value (Qc) (Salem-Garcia et al., 2023). This leaves open the possibility that the activations that we originally associated with confidence in VMPFC actually encode the *sources* of confidence (i.e., value signals) rather than confidence *per se.* To address these concerns, we used the same modelling strategy proposed in Salem-Garcia et al., (2023), and first confirmed their conclusions regarding both learning and confidence models. Indeed, our results showed that the participants choice behavior can be best explained by a reinforcement-learning model featuring context-dependent learning, and confirmatory updating (**Supplementary Figure S1**). Additionally, we did confirm that confidence judgments are best explained by a linear combination of choice difficulty (proxied by the absolute difference between the two available option values) and the chosen option value (Qc) as a biasing term ‒ akin to a choice-congruent evidence integration bias (Miyoshi & Lau, 2020; Peters et al., 2017; Zylberberg et al., 2012). This model provides an excellent fit to participants’ choices and confidence judgments in both learning and transfer phases, and generates key behavioral patterns observed in our data, suggesting that it adequately tracks the cognitive operations mobilized to solve our task. Thereby, the model-derived latent variables allow us to investigate the neural correlate of valuation during learning (Collins & Shenhav, 2022). Note that contrary to most previous studies, our design allowed the separation of option evaluation and motor mapping, which minimizes the potential action-related effect on the correlation between BOLD signal and decision-related variables such as values and confidence (Yoo & Hayden, 2018). In this context, we confirmed that the value of the chosen option correlates positively with BOLD signal in the VMPFC (Baram et al., 2021; Gershman et al., 2009; Gläscher et al., 2009; Skvortsova et al., 2014). More dorsal and lateral regions of the prefrontal cortex (DMPFC, DLPFC) appear to encode with opposite signs the value of the chosen and unchosen options. This pattern could be consistent with the idea that value comparison is effectuated in these more dorsal prefrontal regions (Kolling et al., 2012, 2016; Rangel & Hare, 2010; Wunderlich et al., 2009), and could provide an estimate of the value of control or of information (Klein-Flügge et al., 2022; Shenhav et al., 2016).

To bridge these results on valuation in vmPFC with results suggesting confidence encoding in the same region, we investigated whether VMPFC encode additional confidence precursors (e.g. choice difficulty) in addition to Qc. ROI analyses revealed significant correlations between the activation in the VMPFC and all three confidence precursors identified by our confidence model, suggesting that VMPFC does not simply encode Qc. Consolidating these results, we also found that the activation in the VMPFC can be better explained by confidence than Qc when both variables are included in a single model, and this is observed regardless of the level of granularity considered. Note that, to avoid the double-dipping issue, we selected ROIs that are related to chosen option value from the present study and an independent literature (Bartra et al., 2013), therefore favoring de-facto the opposite hypothesis, namely that VMPFC would preferentially encode Qc. The fact that confidence signals dominate value signals in the VMPFC clashes with the current understanding of its functional role in reinforcement-learning task, which is almost exclusively restricted to option valuation and representation of cognitive maps (Klein-Flügge et al., 2022).

There are at least three tentative explanations for this apparent discrepancy. First, our results could be compatible with the idea that VMPFC does uniquely encode Qc (rather than confidence), but this latent variable is not well estimated by the RL model to robustly capture VMPFC signal variance. In our present modeling exercise as well as a previous modelling paper (Salem-Garcia et al., 2023), we tried to nullify this possibility by going to great length to show that our RL and confidence models can qualitatively and quantitatively account for choice behavior and confidence judgment (**Supplementary Figure S3**). Interestingly, in the (possible) case that a misfit persists and that Qc are mis-estimated, our present results suggest that eliciting confidence judgments could help researchers to better identify the neural networks engaged in value-based learning. Second, similarly to what has recently been shown in decision-making, VMPFC might actually jointly represent decision values and confidence during reinforcement-learning (De Martino et al., 2013; Lebreton et al., 2015a; Lopez-Persem et al., 2020). In our data, only small portions of the VMPFC (anterior and ventral) still correlate positively with Qc when confidence is included in the model. Finally, it is possible that the presence of confidence elicitation in the present study somewhat affects the other computations related to valuation and decision. Although previous work suggests that value and confidence encoding in the VMPFC are both automatic (Lebreton et al., 2009, 2015; Lopez-Persem et al., 2020; Shapiro & Grafton, 2020), an increasing number of studies also reported that VMPFC (value) coding depends on incidental emotional states, as well as specific goals and demands of the task at hand (Cortese, 2021; Engelmann et al., 2015; Sepulveda et al., 2020). These last two possibilities are consistent with the idea that the role of medial and orbital frontal cortex in decision-making and flexible behavior is more complex than initially thought, and might deserve further (re)investigations (Klein-Flügge et al., 2022; Masset et al., 2020).

In the present study, confidence is non-instrumental, and only consists in a read-out of the subjective choice accuracy. In numerous ecological contexts, confidence can be key to monitor and adapt behavioral strategies. Given the multiple layers of confidence and uncertainties uncovered here and the functional dissociations of their neural underpinnings, future studies will need to consider which variable (objective uncertainty, condition-specific confidence, task-wide confidence) and which (confidence) biases impact future behavior – and how. This last point is critical for developing interventions targeting confidence biases, especially as confidence dysfunctions are increasingly seen as relevant markers in clinical applications (Hoven et al., 2019; Rouault, et al., 2022).

## Material and Methods

### Participants

40 participants (female = 23; Age = 22.69±4.44) were recruited from the subject pool of the behavioral science lab (https://www.lab.uva.nl/lab) and through poster adverts distributed on the University of Amsterdam (UvA) campus. The ethical approval was obtained from the Department of Psychology of the Faculty of Social and Behavioral Sciences, at UvA (reference number: 2018-EXT-9205). Before the experiment, only participants that passed the prescreening procedure (e.g., no claustrophobia, no metal in the body) were invited to come to the MRI scanner and were sent an invitation email and detailed information about the experiment and MRI. Participants were asked to arrive at the laboratory 30-min before the experiment. Once participants arrived, they gave informed consent and read the instruction again. Afterward, they experienced a 16-trial practice with the same learning task (but using different symbols) as well as a lottery incentivize procedure outside of the MRI scanner.

The final payout was computed as follows: show-up fee (20€), accumulated outcome from the learning task and bonus from the confidence incentivization procedure. The mean and standard deviation of the payout was 32.18±3.46€. All the tasks were implemented using MatlabR2015a® (MathWorks) and the COGENT toolbox.

### Probabilistic instrumental-learning tasks

We adopted our previous instrumental reinforcement learning task (Lebreton, et al., 2019; Palminteri et al., 2015; Ting et al., 2020) for fMRI by adding incentivized confidence ratings and by separating symbol evaluation and motor response in each trial (see details below). Participants were asked to maximize payoff during the learning task by choosing the symbol with the higher expected value in a pair at each trial (**Figure 1**). In each run of 80 trials, four fixed pairs of abstract symbols were used to represent four conditions in the two (feedback valence: gain or loss) by two (information: partial or complete) within-subjects design (**Figure 1B**). Specifically, eight symbols were divided into four fixed combinations and are constantly arranged to gain & partial (GP), loss & partial (LP), gain & complete (GC), and loss & complete (LC) conditions. Each pair of symbols indicated a specific condition and possible outcomes. For example, for gain contexts (i.e., GP and GC), the possible outcomes are +€1 or +€0.1. Conversely, -€1 or -€0.1 are possible outcomes in the loss contexts (i.e., LP and LC). The probabilistic outcome of an option was determined by reciprocal but independent probabilities, 75% or 25% (**Figure 1B**). The symbol that enjoys a higher expected value (∑ probability × outcome) was defined as the correct option in each pair. Note that only the chosen outcome was added to the final payoff in both the incomplete and complete feedback conditions.

All the participants completed three runs of 80 trials, such that each of the four condition (i.e., each pair of symbols) was repeated 20 times per run. In each trial (**Figure 1A**), the symbols were presented first (1500-3500ms; mean = 2050ms). To avoid the potentially confounding influence of motor responses during symbol evaluation, the symbols disappeared for a while (500-3000ms; mean = 800ms) after symbol presentation. Afterward, two white bars appeared on either right or left of the location of the invisible symbol to indicate which button should be pressed to select the corresponding symbol (i.e., the right button: the white bar was on the right side of the symbol). Once a decision was made, two red bars were displayed beside the chosen symbol (500ms). Before seeing the outcome, participants were asked to state their confidence about choosing the symbol that is better on average (i.e., with a higher expected value). Confidence ratings were done on a scale ranging from 50% to 100% with incremental steps of 5%, and randomized starting points and without time constraints. At the end of each trial, participants were shown the outcome from the chosen option only in the partial information conditions (i.e., GP and LP) for 2000ms. Otherwise, both chosen and unchosen outcomes were displayed in the complete information conditions (i.e., GC and LC. See **Figure 1B**).

In order to motivate participants to accurately report confidence, confidence judgments were incentivized by a Matching Probabilities (MP) mechanism, a well-validated method from behavioral economics adapted from the Becker-DeGroot-Marschak auction (Becker et al., 1964; Ducharme & Donnell, 1973). Specifically, we randomly selected three trials from three runs (i.e., one trial/ run) and then compared the confidence rating *p* at that trial with a random number *r* (chosen from the range between 50% and 100%). If *p* ≥ *r*, then participants won the bonus of 5€ when the chosen symbol indeed had the higher expected value (i.e., the correct one), otherwise, participants won nothing. If *p* < *r*, participants won the bonus of 5€ with a probability of *r*, otherwise, won nothing with a probability of 1-*r*. The euros earned from the game were exchanged for the actual money with a certain exchange rate (1 EU in game = 0.3 payouts EU). Again, all participants were informed about the rule of payout and experienced practice trials in both the learning task and confidence incentivization before the real experiment in the MRI scanner.

### Transfer task

After the learning task, participants left the scanner and were instructed to perform an additional transfer task, where each symbol from the last run of the learning task was paired with all other 7 symbols (i.e., forming 24 new and 4 original pairs). Participants were asked to choose one symbol that can benefit them more, and rate their confidence in their choice. No feedback and monetary incentives were offered in this task. However, participants were asked to imagine that they were able to earn money from the chosen symbols. Because the present study focuses on the neuroimaging data, which was only available for the learning task, analyses of choices from the transfer task are not detailed in the Main Text (but see **Supplementary Figures S2, S4 and S5**).

### Behavioral analyses

In this study, we mostly focused our analyses on three dependent variables of interest available during the learning task: choice accuracy, reaction times, and confidence. The choice accuracy referred to the probability of choosing the relatively better symbol in a pair of symbols (i.e., the one with a higher expected value). The reaction time was defined as the time between the onset of the cues allowing response (referred to as the choice screen in **Figure 1A**) and the actual (self-paced) choice. Confidence simply corresponded to the rating elicited in the confidence judgment screen. To test for the effect of valence and information manipulations, as well as their interaction, these measures were averaged over three runs for each condition and participants and were then fed into two-way repeated-measures ANOVAs. The direction of changes was analyzed by follow-up t-tests. In particular, one-sample t-tests were used when comparing data to a reference value (e.g., guessing level: 50%), and paired t-tests were used to compare responses across different conditions (e.g., gain vs. loss) and different measures (e.g., averaged learning performance vs. averaged confidence).

All statistical analyses were performed using MatlabR2021a® (MathWorks) and its built-in functions (i.e., one-sample t-test: ttest; paired t-test: ttest2; repeated ANOVA: anovan; Pearson’s correlation: corr).

### Computational modelling - methods

#### Learning models – structure and model space

Participants’ choices from both learning task and transfer task were fitted with 10 reinforcement learning models (RL models) proposed in (Salem-Garcia et al., 2023). The models in the model space can be categorized into four families: ABSOLUTE model (ABS), RELATIVE models (REL), ASYMMETRIC models (ASYM), and RELATIVE-ASYMMETRIC models (RELASYM).

The ABS model is the baseline model. Other models were built up based on the ABS model and assumed other sources of information were integrated during learning (**Figure 4A**).

In the ABS model, in all learning contexts *s*, both chosen option value *Q(s, c)* and unchosen option value *Q(s, u)* are updated through a delta-rule function at trials *t*:

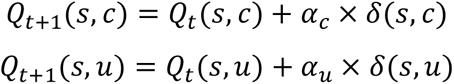

where *α_c_* and *α_u_* are learning rates and δ referred to the prediction error. The prediction error is defined as the difference between the estimated option value Q and the real outcome R:

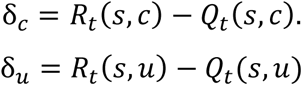

The RELATIVE and RELATIVE-ASYMMETRIC families of models feature context-dependent learning (Bavard et al., 2018; Palminteri et al., 2015; Palminteri & Lebreton, 2021). Thereby, the prediction errors for chosen and unchosen options are corrected with the context value V(s) as follows:

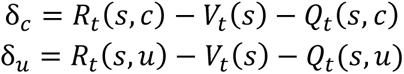

where the context value is also updated through delta-rule with its own learning rate *α_v_* and prediction error *δ_(s, v)_*:

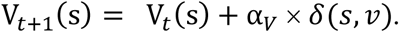

When the counterfactual outcome is available (i.e., complete information conditions), the prediction error for context value is computed as the difference between the estimated context value and the average outcome values:

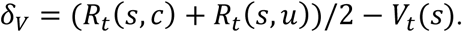

When the outcome for the unchosen option was not available in context s (i.e., partial information conditions), we assume participants infer an approximation of it *Х^∗^*, and calculated the prediction error for context value accordingly:

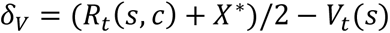

We tested four alternatives for this approximated inference *Х^∗^*, which were implemented in different models. These four alternatives are 0, unchosen option value (*Q_t_*(*s, u*)), the last experienced unchosen outcome for the unchosen option (*R_t−1_*(*s, u*)), and weighted “imaginary forgone outcome” (*w* × ¬*R_t_*(*s*)). Following on our previous work (Salem-Garcia et al., 2023; Ting et al., 2021), the imaginary forgone outcome is determined by the sign of context value (*V_t_*) and the magnitude of the received outcome (*R_t_*(*s, c*)):

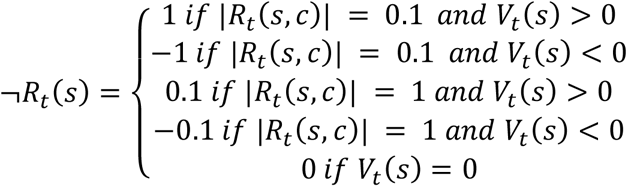

¬*R_t_* is multiplied by a weight parameter *w* (0 ≤ w ≤ 1).

The ASYMMETRIC and RELATIVE-ASYMMETRIC families of models feature asymmetric updating. This follows from previous studies, that demonstrated the presence of a choice-confirmation bias in reinforcement-learning contexts (Lefebvre et al., 2017; Palminteri et al., 2017; Palminteri & Lebreton, 2022). The models capture this bias by allowing two different learning rates (i.e., *α_CON_* and *α_DIS_*) to weight the prediction-error in the value-updating process, depending on the sign of the prediction error. In particular, *α_CON_* (confirmatory learning rate) weights the positive prediction error for chosen option and the negative prediction error for unchosen options. By contrast, α_DIS_ (disconfirmatory learning rate) weights the negative prediction error for chosen options and the positive prediction error for unchosen options.

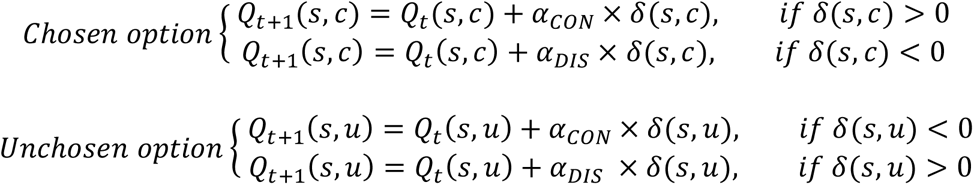

Finally, choice probability between two options (A,B) of the same context s in the learning task is computed with the softmax function:

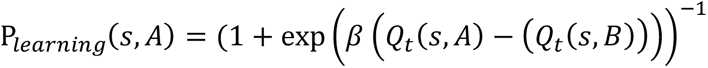

The same softmax function and the same inverse temperature parameter β are applied to model choices in the transfer task between two given options C and D belonging to learning contexts s_C_ and s_D_:

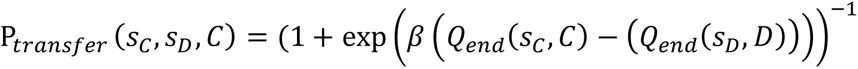

where Q_end_(s_C_,C) and Q_end_(s_D_,D) are the Q-values for options C and D estimated at the end of the learning task in their respective learning contexts.

#### Learning models – model optimization and comparison

Parameter optimization was performed by minimizing the negative logarithm of the posterior probability (n*LPP*) (Daw, 2014):

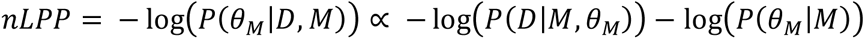

*P*(*D|M, θ_M_*) refers to the likelihood of the observed data *D* (i.e., sequence of choices) given the current model M and its parameters *θ_M_*. *P*(*θ_M_|M*) refers to the prior probability of the parameters.

We used broad priors based on the literature (Daw et al., 2011): The prior distributions of learning rates (α) and weight (w) were defined as beta distributions (Beta(1.1, 1.1) in MATLAB), and the prior distribution of the inverse temperature parameter β was defined as a gamma distribution (Gamma(1.2, 5) in MATLAB). Parameter search was initialized from random starting points selected from certain ranges (i.e., 0 < α < 1; 0 < w< 1; 0 < β < ∞) and used an L-BFGS-B algorithm implemented via Matlab’s *fmincon* function (Byrd et al., 1996).

For model comparison, we calculated, for each individual, the Laplace approximation to the model evidence (LAME), which penalizes model’s complexity (i.e., number of parameters) as follow:

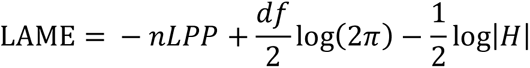

where n is the number of trials, *df* is the number of free parameters and *H* is the determinant of the Hessian.

Quantitative model comparison was performed via a formal Bayesian Model Selection (BMS) random-effect procedure, as described in (Daunizeau et al., 2014) and implemented in the mbb-vb-toolbox (http://mbb-team.github.io/VBA-toolbox/). This toolbox performs the Bayesian model selection procedure and estimates two indicators: the expected frequencies (*EF*) and the exceedance probability (EP) for each model. Specifically, the expected frequency *EF* of a model quantifies the probability that the model generated the data for any randomly selected subject. Note that the EF should be higher than chance level given number of models in the model space. Exceedance probability (EP), on the other hand, quantified the belief that the model is more likely than all the other models of the model-space.

Note that parameter recovery and model recovery for the learning models are detailed in (Salem-Garcia et al., 2023).

#### Confidence models – structure and model space

Participants’ confidence ratings were separately fitted in the learning task and in the transfer task with four confidence models proposed in (Salem-Garcia et al., 2023). Confidence models are defined as logit-transformed multiple linear regression models that use the latent variables estimated by the winning RL model (i.e., RELASYM) to predict confidence ratings (**Figure 5A**). Each model consists of one intercept and two predictors: (1) task difficulty, which is measured as absolute value difference between options (|Qc-Qu|) and (2) a hypothesized source of valence bias. We tested four hypothesized sources of valence bias: none (0), the summed value of available options (ΣQ = Qc + Qu), the expected value of the chosen option (Qc), and the context value (V). In the learning task, this latter was straightforwardly available as V_t_(s). In the transfer task, we generalized the idea of context value for choice between any two options C and D, as 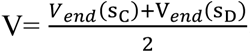, where V_end_(s_C_) and V_end_(s_D_) are the (choice-independent) values associated with the original contexts of options C and D estimated at the end of the learning task. In addition to these two predictors, the models for the learning task contains an additional predictor capturing the fact that confidence in the current trial are usually influenced by confidence in the previous trial: an autocorrelation term Conf_t-1_. Ultimately, confidence models can be expressed as followed:

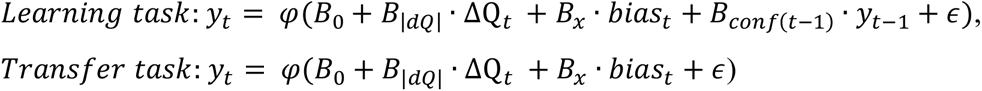

where y refers to confidence ratings, bias can be either 0, ΣQ, Qc or V in different models, and *ε* is the error term (sampled from a Gaussian distribution with zero mean). φ(x) is the logistic link function φ(x) = 1/(1 + e^−x^).

#### Confidence models ‒ model optimization and comparison

Confidence model parameters were estimated by fitting robust linear regression, via the procedure of maximizing log-likelihood (LL), as implemented in MATLAB robustfit functions. Considering that no principled priors for the confidence models are available, we used LL to approximate model evidence for each subject and each model as the BIC (Bayesian information criterion), defined as

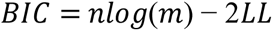

where n is the number of parameters and m is the number of data points (trials). Similarly to the learning models, we fed the BIC (from each subject in each model) to the random-effect BMS routine implemented in the mbb-vb-toolbox (http://mbb-team.github.io/VBA-toolbox/; Daunizeau et al., 2014).

Note that parameter recovery and model recovery for the confidence models are also detailed in (Salem-Garcia et al., 2023).

### fMRI

#### fMRI acquisition

The fMRI data were acquired using a 3.0-Tesla Philip Achieva scanner with 32 channels head array coil. We recorded both structural images and functional brain images. T1 weighted structural scans were recorded with the following parameters: FOV (Field of View): 240×180×220 mm^3^, Voxel size =1×1×1 mm^3^, TR = 8.2ms and TE = 3.7ms. Each T2*-weighted functional scan consisted of 36 axial echo-planar images (EPI) acquired in ascending sequence with voxel size of 3×3×3 mm^3^, slice gap = 0.3 *mm*, TR= 2000ms, TE = 28ms and the flip angle of 76°. Each subject completed 3 runs in a scanning session. Given the task was self-paced and the fMRI scanner was manually terminated (i.e., ∼10 seconds after the last feedback phase), the total numbers of functional scans for each subject in each run were not the same. Most participants completed the task in around 15 minutes. The field maps (i.e., magnetic field’s inhomogeneity) were collected as well between the second and the third run.

#### fMRI preprocessing

The functional images were preprocessed using SPM12 (Wellcome Department of Imaging Neuroscience, London) with the following steps: realignment and unwarp, co-registration, segmenting anatomical images, normalization, and smoothing. To correct for potential head movement during functional images collection, all functional volumes (from three runs) were realigned to the first volume in the first run and were un-warped with collected field maps. To improve the quality of the following normalization, the mean functional (the output from realignment) and anatomical images were co-registered. The anatomical image from each subject was segmented into six images (i.e., grey matter, white matter, cerebrospinal fluid, fat tissue and air) using nonlinear deformation fields and SPM12’s Tissue Probability Maps (TPMs). All segmented images were then normalized to the Montreal Neurological Institute T1 template (i.e., MNI152) using forward deformation fields from the segmentation output. Finally, the EPI images were normalized and smoothed with a full width half maximum Gaussian kernel of 6-mm (2 times of voxel size of functional images) full-width at half maximum (FWHM) isotropic Gaussian kernel.

#### fMRI analysis: GLMs

Our fMRI analyses leveraged a total of 5 different GLMs (whose specificities are briefly described below, and summarized in **Table 1**). All GLMs modelled separately the four main events composing our prototypical trial: symbol presentation, choice, confidence rating, outcome. These event-related regressors were modeled using boxcar functions with corresponding durations. Across all models, the choice and confidence onsets were respectively modulated with parametric modulators accounting for (1) choice (right or left), (2) the distance between initial and final rating point for rating onset. Across all models, all parametric modulators were z-scored to ensure results from different conditions and regressors were comparable (Lebreton, et al., 2019). To allow different regressors to fairly compete in explaining the same share of data variance, SPM serial orthogonalization was turned off. To remove motion artifact and to improve the quality of fMRI results, all the GLMs also contained six realignment parameters, which were created during preprocessing. Linear contrasts of regression coefficients were designed at the individual level (first-level), and, unless otherwise specified, taken to the group-level random-effect analysis (second-level). For whole brain analyses, second-level analyses consisted in one-sample t-test, whose statistical significance was defined with whole-brain cluster-defining height threshold at uncorrected *p*<.001 and family-wise error (FWE)-corrected threshold of *p*<.05. For ROI analyses, the individual-level averaged contrast values were extracted from the ROI using spm built-in function (i.e., spm_get_data.m). These values were than taken to second-level analyses, consisting in one-sample or paired t-tests, as well as two-way repeated-measures ANOVAs.

GLM1 divided symbol onset and outcome onset into four conditions each (i.e., GP, LP, GC, LC). These eight events of interest were enriched with parametric modulators accounting for 1) confidence ratings for each condition-specific symbol onset, 2) received outcome (coded as 1/ 0 for a relatively good /bad outcome) for each condition-specific outcome phase.

GLM2_WID_ and GLM2_SPE_ featured a single regressor for the symbol and for the outcome events, effectively concatenating all conditions. GLM2_WID_ and GLM2_SPE_ only differed from each other regarding the variable used as the confidence parametric modulator. In GLM2_WID_, confidence consisted in the native ratings. In GLM2_SPE_, confidence ratings were first z-scored *per condition* and before being re-concatenated as a single variable.

GLM3-5 implemented model-based fMRI, and leveraged the latent variables obtained from our winning computational model see **Methods and Figure 4A** (see also **Figure S1)**.Because the computational variables are meant to capture the difference between conditions, these GLMs also featured a single regressor for the symbol and for the outcome events. As is customary in functional neuroimaging studies, and although beyond the scope of this manuscript, all those GLMS featured the modelled prediction-error (PE) as a parametric modulator of the outcome event.

In GLM3, the symbol presentation onset was modulated by Qc (chosen option value), Qu (unchosen option value), and V (context value).

In GLM4, the symbol presentation onset was modulated by Qc (chosen option value), |Qc-Qu| (absolute value differences), and previous confidence (conf_t-1_).

In GLM5, the symbol presentation onset was modulated by confidence and Qc (chosen option value.

#### ROI analyses

ROIs were created using the marsbar toolbox (Brett et al., 2002). A first family of ROIs was built from the significant clusters from the GLM1 confidence activations (VMPFC, dmPFC, Inferior Frontal Gyrus, and Insula)

Alternative VMPFC ROIs were also built from independent meta-analyses (Bartra et al., 2013) and from significant clusters from other analyses of the resent study (e.g., voxels significantly correlated to Qc in GLM3).

#### Bayesian model selection (fMRI)

Bayesian model selection (BMS) was effectuated using SPM’s toolbox: MACS (Soch & Allefeld, 2018). In the first step (i.e., model assessment), the first-level GLMs of interest from each subject were used to estimate voxel-wise cross-validated log model evidence (cvLME) maps. The maps were generated for each GLM and each subject within the model space. In the second step (i.e., model comparison and selection), the cvLME maps served as inputs for the cross-validated Bayesian Model Selection (cvBMS) to compare GLMs within the model space. Only voxels available in all participants were included in those analyses.

## Acknowledgments and fundings

The authors thank Tiffany M. Hrkalović for assistance with the fMRI data collection. This study was funded by startup funds from the Amsterdam School of Economics awarded to JBE. CCT is supported by GSSA, MOE Taiwan Scholarship (1081007012). ML is supported by an ERC Starting Grant (INFORL-948671). SP is supported by an ERC Consolidator Grant (RaReMem-101043804) and by the Agence National de la Recherche (CogFinAgent: ANR-21-CE23-0002-02; RELATIVE: ANR-21-CE37-0008-01; RANGE: ANR-21-CE28-0024-01)

## Authors contributions

All authors designed the study. JBE acquired funding. CCT ran the experiment under the guidance of JBE. CCT conducted the analysis, and drafted the manuscript under the supervision of ML. NSG performed the model-validation analyses. All authors critically assessed and discussed the results, and revised and approved the manuscript.

## Notes

### Competing Interest Statement

The authors have declared no competing interest.

