## Supplementary Materials for "Neural and computational underpinnings of biased confidence in human reinforcement learning"

#### Table of Contents

*Supplementary Results* .......... *2*

**Computational modelling - methods** .......... Error! Bookmark not defined.

Learning models – structure and model space .......... **Error! Bookmark not defined.**
Learning models – model optimization and comparison .......... **Error! Bookmark not defined.**
Confidence models – structure and model space .......... **Error! Bookmark not defined.**
Confidence models – model optimization and comparison .......... **Error! Bookmark not defined.**

### Supplementary Results

#### Computational modelling - results

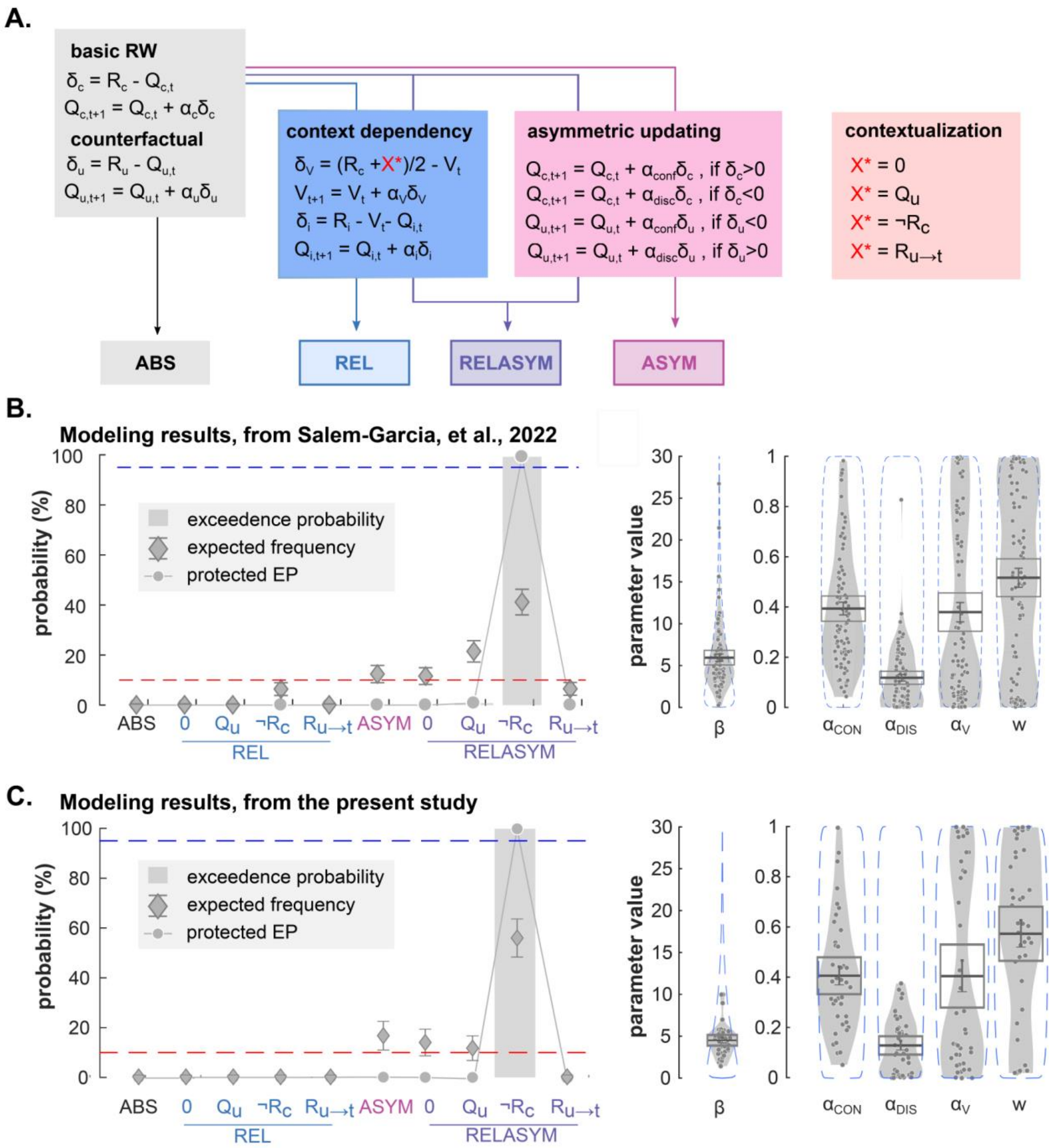

**Figure S1 | Modeling approach and results. (A)** The model architecture and family Bayesian Model Comparison (BMC) procedure. Color panels represent different components of value updating rules. Gray panel: Absolute model (ABS), which consists of basic Rescorla-Wagner (RW) update rule. This rule updates chosen and unchosen option values via outcome directly. Blue panel: Relative model (REL), which consists context-dependent component and updates option values by considering context value. Pink panel: Asymmetric updating model (ASYM), which updates option values based on the valence of prediction error. Purple panel: relative-asymmetric model (RELASYM), which is the combination of relative model and asymmetric updating model. The contextualization panel is used to update unchosen option in the partial information condition. Specifically, X* is determined as the unchosen option outcome (Ru) when the value is available in the complete information condition. When the unchosen option outcome (Ru) is not available in the partial information condition, X* is hypothesized as none (0), expected unchosen value (Qu), paired outcome (¬R) and last seen outcome associated with the option (Ru→t). **(B)** Modeling results from Salem-Garcia et al. 2023 and **(C)** from the present study. **(B-C)** Left panels: Bayesian model comparison. X-axis represents the models with different hypothesized contextualization values. Y-axis represents the value of three criteria, including exceedance probability (grey histograms), expected frequencies (diamonds) and protected exceedance probability (line & dots) of each model. The red dashed line represents the guessing level for EF and LF. The blue dashed line represents the threshold (95%) for the exceedance probability. **(B-C)** Right panels: Estimated parameter values of the winning model (ASYMREL, X* = with ¬Rc) from Salem-Garcia et al. 2023 **(B)** and from the present study **(C).** Dots represent individual data points. Error bars displayed within the violin plots indicate the sample mean ± SEM. The blue, dotted envelop represent the prior distribution.

$Q_{c/u,t}$: value of the chosen/unchosen option at trial t. $R_{c/u}$: outcome associated to the chosen/unchosen option. $\delta_{c/u}$: prediction error for the chosen/unchosen option. $\alpha_{u/c}$: learning rate for the chosen/unchosen option. $\alpha_{conf/disc}$: learning rate for confirmatory/disconfirmatory information. $V_t$: context value; $\delta_{V:}$ prediction error for the context value. $\alpha_V$: learning rate for the context value.

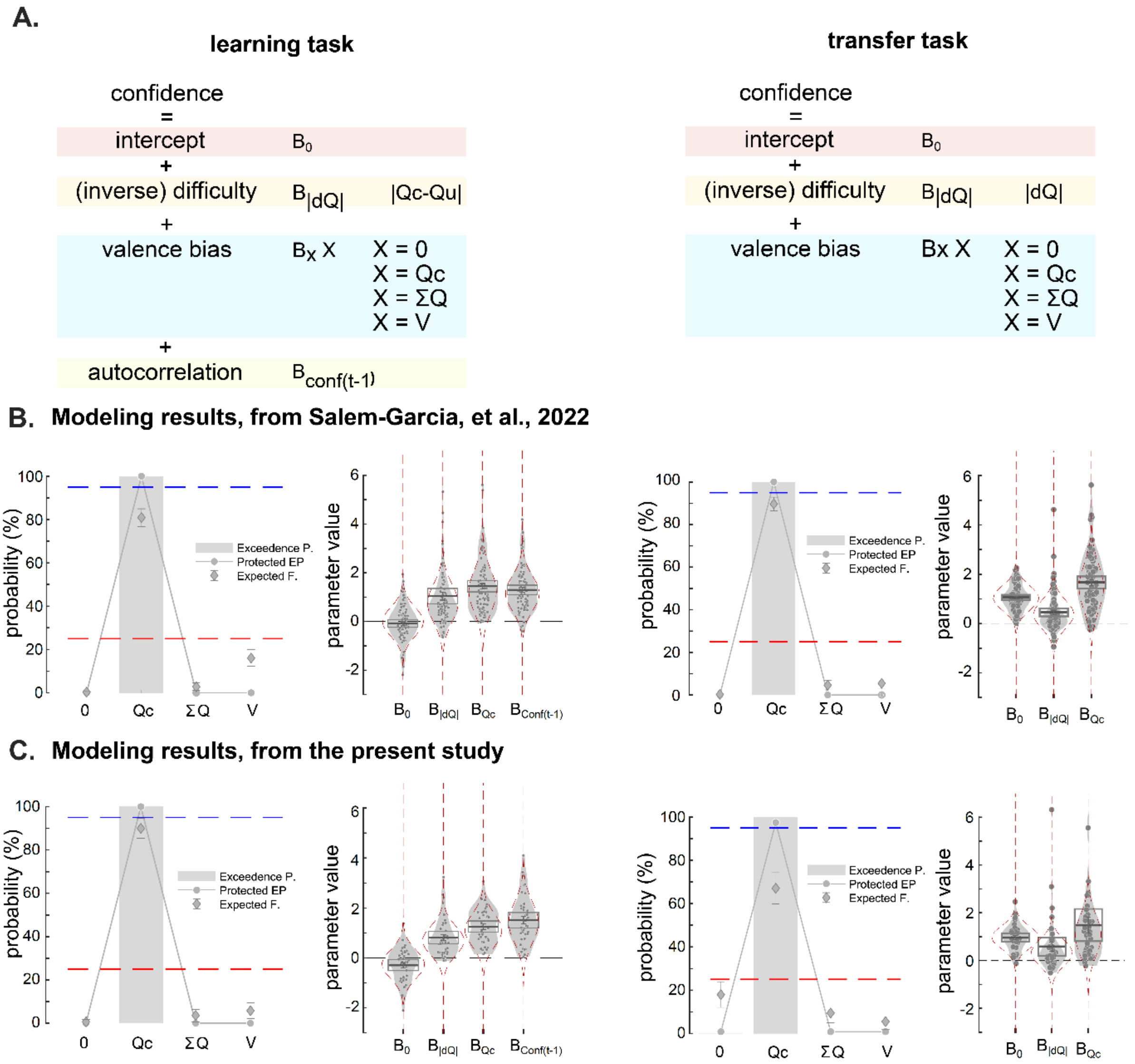

**FigureS2 | Modeling confidence in the learning task and transfer task. (A)** Confidence models are linear regression models. *Left:* Each model in the learning task consists of an intercept and three slopes, which accounts for difficulty (i.e., absolute value difference of present options), source of valence bias, and autocorrelation (i.e., confidence ratings in the previous trial). *Right:* Each model in the transfer task consists of an intercept and two slopes, which accounts for difficulty, and source of valence bias. In different models, the source of valence bias (X) is hypothesized as none, or summed value of present options, or context value, or expected chosen option value. These values are estimated by the winning RL model. **(B)** Modeling results from Salem-Garcia et al. 2023 and **(C)** from the present study. **(B-C)** *Left panels* of the learning task and the transfer task: Bayesian model comparison. X-axis represents the models with different hypothesized source of confidence bias. Y-axis represents the value of three criteria, including exceedance probability (grey histograms), expected frequencies (diamonds) and protected exceedance probability (line & dots) of each model. The red dashed line represents the guessing level for EF and LF. The blue dashed line represents the threshold (95%) for the exceedance probability. **(B-C)** *Right panels* of the learning task and the transfer task: Estimated regression coefficients of the winning model (with valence bias =Qc) from Salem-Garcia et al. 2022 **(B)** and from the present study **(C).** Dots represent individual data points. Error bars displayed within the violin plots indicate the sample mean ± SEM. The red, dotted envelop represent the prior distribution.

Qc: chosen option value; ΣQ: summed present option values; V: context value; β: inverse temperature; $\alpha_{CON/DIS}$: learning rate for confirmatory/disconfirmatory information; $\alpha_V$: learning rate for the context value; w: partial contextualization weight.

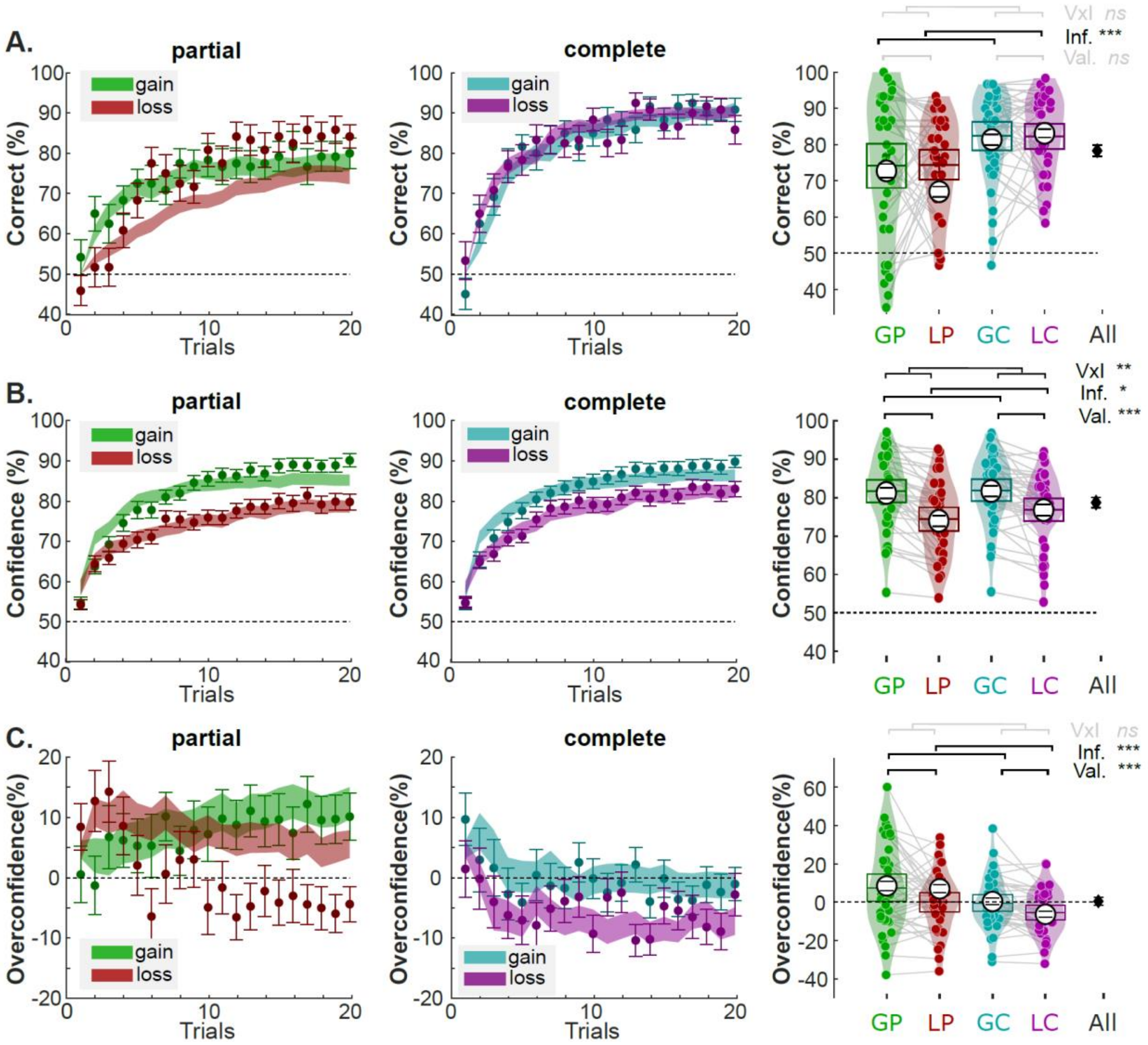

**FigureS3 | Model predictions and real data in the learning task (A-C).** Left and middle panels are trial-by-trial **(A)** percentage of correct responses, **(B)** Confidence rating, and **(C)** overconfidence in the partial information (left panels) and complete information condition (middle panels). Filled colored areas represent mean±SEM. Shaded areas represent the model predictions. **(A-C)** Right panels are average **(A)** percentage of correct responses, **(B)** Confidence rating, and **(C)** overconfidence across conditions at the individual level (dots) and group-level (horizontal bars). The black error bars indicate the overall performance over conditions. White circles and error bars displayed within the violin plots indicate mean ± SEM of the model predictions.

G75 and G25: options associated with 75% and 25% probability of winning the big gain (1€), respectively; L75 and L25: options associated with 75% and 25% probability of losing the big loss (-1€), respectively.

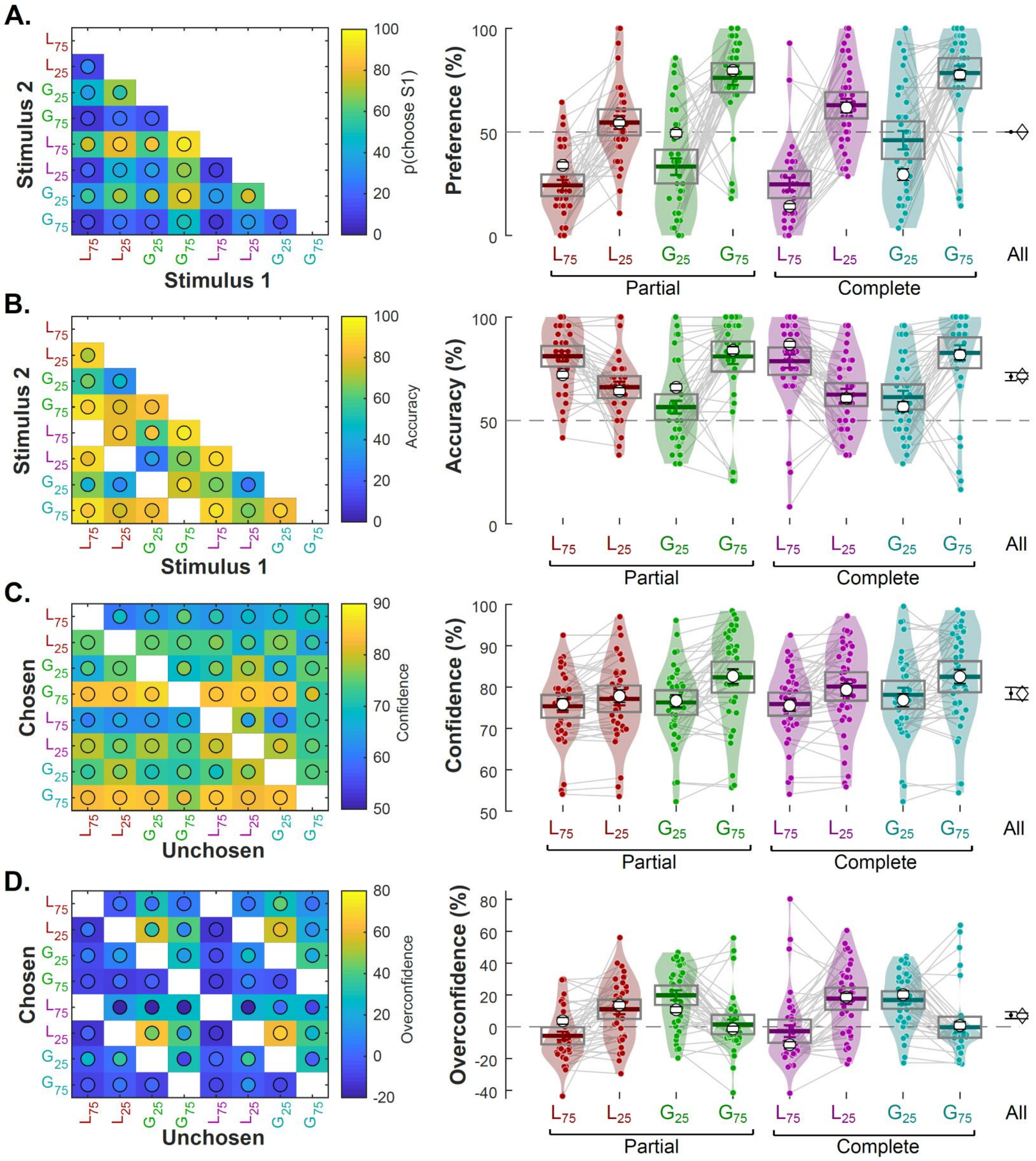

**Figure S4 | Model predictions and real data in the transfer task. (A-D)** Left: Color-coded preference (A), accuracy (B), confidence (C) and overconfidence (D) for each pair of option presented in the transfer task. Lego plot pictures color-coded confidence for each pair of option presented in the transfer task. Background (square) color represent behavior, while overlaid colored circles represent model predictions. Right: Individual averaged confidence for each option (i.e. averaged over every choice including the option). Connected dots represent individual data points in the within-subject design. Error bars displayed within the violin plots indicate the sample mean ± sem. Rightmost black diamond and error bar represents the average over all conditions (mean ± sem). White circles and error bars represent mean ± sem of the model predictions.

G75 and G25: options associated with 75% and 25% probability of winning the big gain, respectively; L75 and L25: options associated with 75% and 25% probability of losing the big loss, respectively

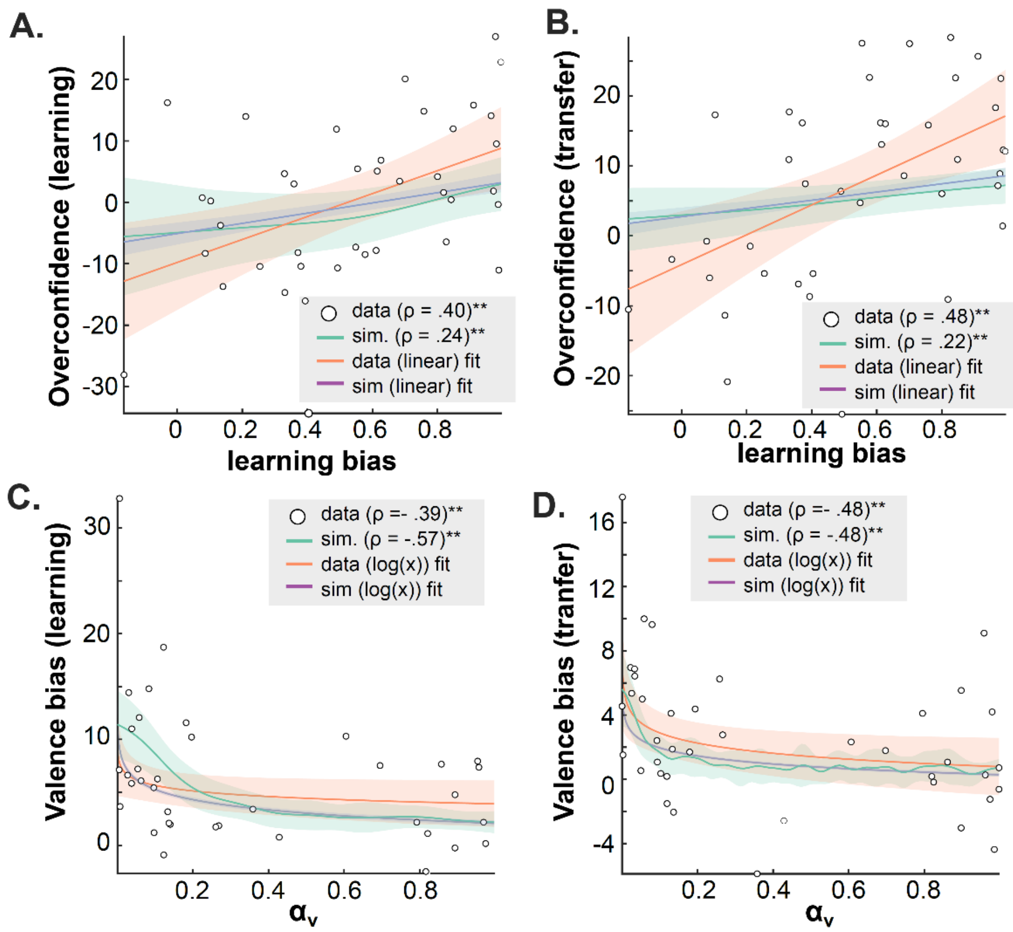

**Figure S5 | Correlations between learning model parameters and confidence biases.** The scatter-plots depict correlations estimated in both the learning (A, C) and the transfer (B, D) tasks, between learning bias ((αCON - αDIS)/(αCON + αDIS)) and overconfidence (A, B) and between contextual learning rate (αV) and valence-induced confidence bias (C, D). Dots represent individual estimates. Green lines and shaded areas represent non-parametric regression estimate ± CI 95%, obtained from simulations (see Methods).

### fMRI – supplementary results

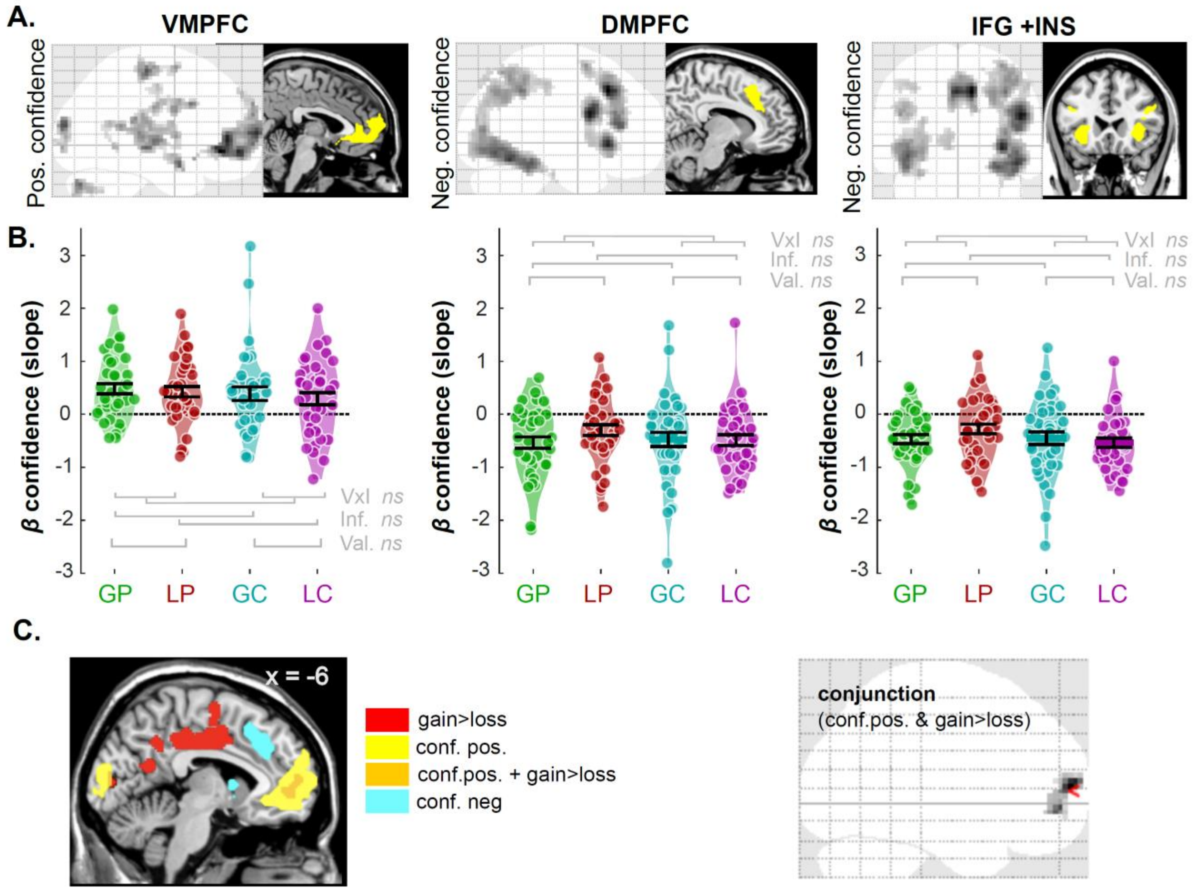

**Figure S6 | confidence encoding in the human brain. (A)** Results of GLM1 whole-brain analysis, reproduced from **Figure 3A**, for convenience. Brain areas positively (left panels) and negatively (middle and right panels) correlate with confidence rating during the symbol presentation phase. Significant voxels are displayed on the glass brains in a gray-to-black gradient manner ($p_{uncorrect}$< .001, cluster size >47). The yellow areas in the anatomical brain are ROIs (vmPFC, dmPFC, and IFG+INS), which are used in the following ROI analyses. **(B)** Violin plots represent the sample distribution of fMRI regression coefficients of parametric confidence signals for the different contexts (represented by different colors), in the ROI depicted in (A). Dots correspond to individual regression coefficients. Error bars represent sample mean ± SEM. GP: gain/partial; LP: loss/partial; GC: gain/complete; LC: loss/complete. **(C)** Left: superposition of GLM1 significant clusters, for 3 contrasts: [gain>loss], red; [positive effect of parametric confidence], yellow; [negative effect of parametric confidence]. Only the first two overlapped (orange). Significant clusters defined as: voxel-wise $P_{uncorrected}$ < .001; cluster-wise $P_{FWE}$ < .05. Right: formal whole brain analysis of a conjunction between [positive effect of parametric confidence] and [gain vs loss], (voxel-wise $P_{uncorrected}$ < .001; cluster-wise $P_{FWE}$ < .05).

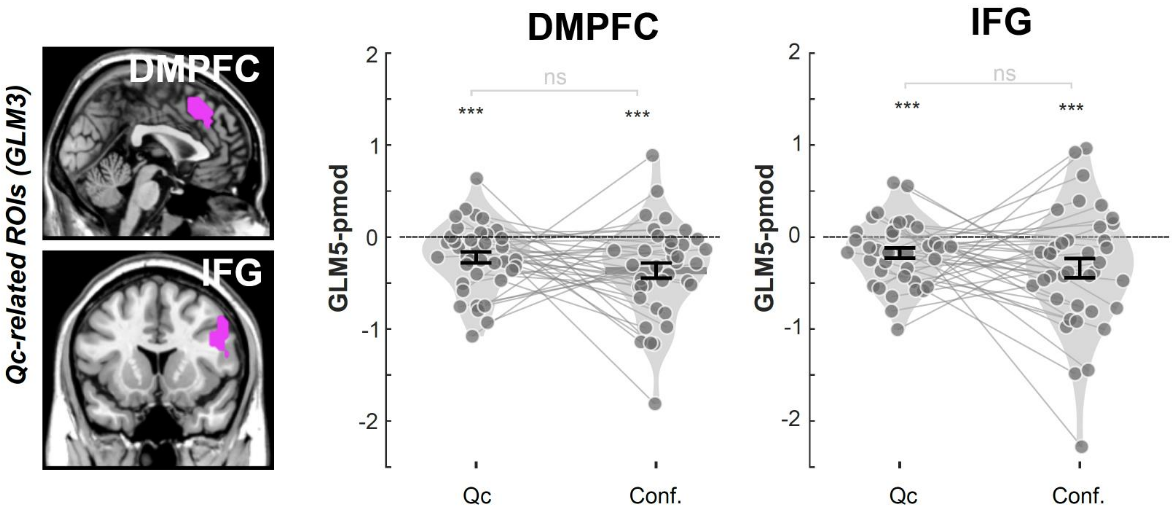

**Figure S7 | Activations in the negative networks cannot be better explained by confidence. (A)** ROI analysis with Qc-related ROIs identified in the present study (purple areas). **(B)** The regression coefficients corresponding to Qc and confidence in GLM5 were summarized at the individual level (dots) for ROI of DMPFC and IFG+INS, separately. Error bar represents SEM. **Qc**: parametric modulator of chosen option; **Conf.**: parametric modulator of confidence ratings;

~: .05<*p*<.1; *: .001<*p*<.01; **: .01<*p*<.001; ***: *p*<.001

### Supplementary tables

**A.**

| **Gender** M/F | **Age** mean ± STD | **Performance (%)** mean ± SEM | **Confidence (%)** mean ± SEM | **RT (ms)** mean ± SEM | **Conf. RT** mean ± SEM |
|---|---|---|---|---|---|
| 17/23 | 22.69±4.44 | 78.31 ± 1.59 | 78.71 ± 1.33 | 715.76 ± 33.76 | 1948±68.13 |

**B.**

| | Gain Partial | Loss Partial | Gain Complete | Loss Complete |
|---|---|---|---|---|
| **Performance (%)** mean ± SEM | 74.12 ±0.48 | 74.42 ± 0.32 | 82.42 ± 0.31 | 82.29 ± 0.27 |
| **Confidence (%)** mean ± SEM | 81.63 ± 0.23 | 74.38 ± 0.24 | 81.92 ± 0.23 | 76.89 ± 0.23 |
| **Confidence bias** mean ± SEM | 7.51 ± 3.50 | -0.04 ± 2.45 | -0.49 ± 2.20 | -5.41 ± 3.50 |
| **RT (ms)** mean ± SEM | 693.02 ± 34.02 | 745.91 ± 38.02 | 698.31 ± 34.34 | 725.79 ± 38.20 |
| **Conf. RT (ms)** mean ± SEM | 1960±66.58 | 1909±68.38 | 1980±74.42 | 1943±74.04 |

**Table S1. Demographics and descriptive statistical results of behavioral data**
M: Male; F: Female; STD: standard deviation; SEM: standard error of the mean

| | Valence | Information | Valence × Information |
|---|---|---|---|
| **Performance**<br>F(1,39) [$\eta^2$] (*p*-val.) | 0.00 [0.00]<br>(.9666) | 22.05 [0.07]<br>(<.0010)*** | 0.01 [0.00]<br>(.9056) |
| **Confidence**<br>F(1,39) [$\eta^2$] (*p*-val.) | 36.56 [0.10]<br>(<.0010)*** | 6.76 [0.00]<br>(.0131)* | 9.62 [0.00]<br>(.0036)** |
| **Confidence bias**<br>F(1,39) [$\eta^2$] (*p*-val.) | 12.28 [0.03]<br>(.0012)** | 14.42 [0.04]<br>(<.0010)*** | 0.58 [0.00]<br>(.4506) |
| **RT**<br>F(1,39) [$\eta^2$] (*p*-val.) | 4.77 [0.00]<br>(.0350)* | 0.31 [0.00]<br>(.5782) | 0.97 [0.00]<br>(.3318) |
| **Conf. RT**<br>F(1,39) [$\eta^2$] (*p*-val.) | 3.42 [0.00]<br>(.0722)~ | 1.62 [0.00]<br>(.2112) | 0.09 [0.00]<br>(.7691) |

**Table S2. Repeated measures ANOVA results reported for learning performance**

Valence: gain/loss; Information: partial/complete. ~ *p*<.10; * *p* <.05; ** *p* <.01; *** *p* <.001

| | | GLME1 | GLME2 | GLME3 | GLME4 |
|---|---|---|---|---|---|
| **Valence** | β±SE<br>t-val<br>(*p*-val) | 4.95±1.06<br>4.66<br>(<.001)*** | 2.55±0.55<br>4.61<br>(<.001)*** | 5.57±1.19<br>4.66<br>(<.001)*** | 2.96±0.63<br>4.67<br>(<.001)*** |
| **Information** | β±SE<br>t-val<br>(*p*-val) | -2.44±0.67<br>-3.60<br>(<.001)*** | -1.16±0.36<br>-3.21<br>(.001)** | -2.27±0.75<br>-2.99<br>(.002)** | -1.27±0.40<br>-3.14<br>(.001)** |
| **Valence × Information** | β±SE<br>t-val<br>(*p*-val) | 2.13±0.70<br>3.02<br>(<.001)*** | 1.03±0.43<br>2.38<br>(.017)* | 1.83±0.83<br>2.19<br>(.028)* | 1.10±0.43<br>2.51<br>(.011)** |
| **RT** | β±SE<br>t-val<br>(*p*-val) | -0.00±0.00<br>-13.94<br>(<.001)*** | -0.00±0.00<br>-11.65<br>(<.001)*** | -0.00±0.00<br>-13.22<br>(<.001)*** | -0.00±0.00<br>-11.62<br>(<.001)*** |
| **Session** | β±SE<br>t-val<br>(*p*-val) | | 0.52±0.24<br>2.17<br>(.029)* | 0.61±0.21<br>2.18<br>(.002)** | 0.42±0.20<br>2.02<br>(.042)* |
| **Previous confidence (regardless of conditions)** | β±SE<br>t-val<br>(*p*-val) | | | 0.42±0.00<br>50.32<br>(<.001)*** | 0.12±0.00<br>14.16<br>(<.001)*** |
| **Previous confidence (Within condition)** | β±SE<br>t-val<br>(*p*-val) | | 0.55±0.00<br>72.31<br>(<.001)*** | | 0.49±0.00<br>57.40<br>(<.001)*** |
| **$R^2$** | | 0.3260 | 0.6156 | 0.4970 | **0.6238** |

**Table S3. Estimated coefficients from generalized linear mixed-effect models (GLME) on confidence**

β: estimated regression coefficients for fixed effects. SE: estimated standard error of the regression coefficients.

~ $p$ <.1; * $p$ <.05; ** $p$ <.01; *** $p$ <.001

| | Gain partial | Loss partial | Gain complete | Loss complete | Overall |
|---|---|---|---|---|---|
| mean ± SEM | -0.18 ± 0.03 | -0.21 ± 0.03 | -0.18 ± 0.02 | -0.20 ± 0.03 | -0.19 ± 0.02 |
| t(39) | -5.56 | -6.13 | -6.63 | -5.86 | -8.21 |
| (*p*-val) | (<.001)*** | (<.001)*** | (<.001)*** | (<.001)*** | (<.001)*** |

**Table S4. Correlation between confidence and RT**

The correlation between confidence and reaction time was performed at the session level using Pearson's R, then averaged at the individual level. Reported statistics correspond to a random-effects analysis (one sample t-test) performed at the population level.

**SEM**: standard error of the mean. **t**: Student t-value.

~ *p*<.10; * *p* <.05; ** *p* <.01; *** *p* <.001

| | | Exp. 1 | Exp. 2 | Exp. 3 | Exp. 4 | Exp. 5 | Exp. 6 | **fMRI** |
|---|---|---|---|---|---|---|---|---|
| **Intercept** | β ± SE | 8.02 ± 2.15 | 2.94 ± 1.29 | 2.16 ± 0.95 | 3.54 ± 1.37 | 6.76 ± 1.81 | 3.59 ± 1.95 | 5.02 ± 0.84 |
| | t-val | 3.72 | 2.27 | 2.27 | 2.58 | 3.73 | 2.41 | 5.97 |
| | (*p*-val) | (.002)** | (.037)* | (.038)* | (.020)* | (.002)** | (.028)* | (<.001)*** |
| **Slope** | β ± SE | -0.00±0.01 | -0.03 ± 0.01 | -0.02 ± 0.04 | -0.06 ± 0.06 | 0.17 ± 0.18 | -0.10 ± 0.03 | -0.01± 0.01 |
| | t-val | -0.27 | -3.55 | -0.46 | -0.97 | 0.81 | -3.35 | -0.97 |
| | (*p*-val) | (.793) | (.003)** | (.662) | (.35) | (.368) | (.004)** | (.339) |

**Table S5. Comparison of Ting et al., 2020 and current estimated coefficients from inter-individual robust regressions.**

For each individual, we estimated the net effect of valence on RT and confidence, by computing the averaged difference of these behavioral measures in the gain versus loss contexts. The fMRI experiment with larger sample size (n=40; selected by a orange frame) revealed the similar results as we found from out previous result : Experiment 1-6 in (Ting et al., 2020).

β: estimated regression coefficient. SE: estimated standard error of the regression coefficient.

~ *p*<.10; * *p* <.05; ** *p* <.01; *** *p* <.001

| *cue-evoked* | | Valence | Information | Valence × Information |
|---|---|---|---|---|
| **VMPFC** | F(1,37) [$\eta^2$] (*p*-val.) | 8.99 [0.020] (.0048)** | 0.34 [0.000] (.5602) | 3.99 [0.008] (.0532) |
| **DMPFC** | F(1,37) [$\eta^2$] (*p*-val.) | 3.05 [0.007] (.0893) | 0.30 [0.000] (.5849) | 0.22 [0.000] (.6449) |
| **IFG+INS** | F(1,37) [$\eta^2$] (*p*-val.) | 0.13 [0.000] (.7119) | 0.03 [0.00] (.8721) | 0.04 [0.00] (.8421) |
| ***Pmod: confidence*** | | Valence | Information | Valence × Information |
| **VMPFC** | F(1,37) [$\eta^2$] (*p*-val.) | 0.36 [0.003] (.5502) | 1.56 [0.006] (.2194) | 0.03 [0.000] (.8429) |
| **DMPFC** | F(1,37) [$\eta^2$] (*p*-val.) | 1.23 [0.006] (.2752) | 1.13 [0.002] (.2938) | 0.97 [0.008] (.3317) |
| **IFG+INS** | F(1,37) [$\eta^2$] (*p*-val.) | 0.34 [0.002] (.5629) | 4.26 [0.010] (.0459) | 2.41 [0.013] (.1287) |

**Table S6. Repeated measures ANOVA results reported for ROI analysis (GLM1 during symbol presentation)**

~ $p<.10$; * $p<.05$; ** $p<.01$; *** $p<.001$ with Bonferroni corrected for 3 comparisons.

| Parametric effect | Regions | MNI coordinates | | | Cluster size | T |
|---|---|---|---|---|---|---|
| | | x | y | z | | |
| ***Confidence (Positive effect)*** | pregenual anterior cingulate cortex (pgACC) | 3 | 35 | -7 | 697 | 6.65 |
| | | 0 | 41 | 2 | | 6.11 |
| | | -9 | 50 | -7 | | 6.00 |
| | Calcarine sulcus | -6 | -91 | 14 | 79 | 5.42 |
| | | -3 | -94 | -4 | | 4.23 |
| | | -12 | -91 | 5 | | 4.18 |
| | Precentral gyrus | -30 | -25 | 59 | 139 | 5.37 |
| | | -18 | -31 | 62 | | 3.88 |
| | | -45 | -25 | 56 | | 3.58 |
| | Middle temporal gyrus | -57 | -25 | -4 | 65 | 5.21 |
| | | -48 | -31 | 2 | | 4.22 |
| | Middle temporal gyrus | 60 | 2 | -19 | 397 | 5.18 |
| | | 66 | -19 | 2 | | 4.94 |
| | | 69 | -31 | 8 | | 4.68 |
| | Cerebellum | -27 | -79 | -34 | 68 | 5.07 |
| | | -21 | -85 | -43 | | 3.90 |
| | Middle frontal gyrus | 24 | 11 | 32 | 130 | 4.87 |
| | | 27 | -1 | 26 | | 4.57 |
| | | 21 | -19 | 29 | | 4.46 |
| | Cerebellum | 27 | -79 | -34 | 86 | 4.83 |
| | | 42 | -70 | -40 | | 4.11 |
| | | 45 | -61 | -40 | | 3.72 |
| | Superior temporal gyrus | -51 | -34 | 20 | 73 | 4.60 |
| | | -63 | -46 | 14 | | 4.54 |
| | | -60 | -55 | 17 | | 4.09 |
| | Precentral gyrus | 21 | -31 | 59 | 58 | 4.51 |
| | | 30 | -28 | 62 | | 4.22 |
| | Frontal gyrus | 15 | -19 | 62 | 61 | 4.50 |
| | | 6 | -1 | 62 | | 4.48 |
| | | 21 | -19 | 56 | | 4.05 |
| ***Confidence (Negative effect)*** | Dorsal medial prefrontal cortex (DMPFC) | 9 | 23 | 41 | 324 | 8.97 |
| | | -3 | 26 | 41 | | 7.83 |
| | | 0 | 23 | 53 | | 5.15 |
| | Inferior frontal gyrus (IFG) | 48 | 8 | 23 | 405 | 8.41 |
| | | 45 | 32 | 14 | | 6.62 |
| | | 45 | 29 | 23 | | 5.87 |
| | Insula (INS) | 33 | 23 | -4 | 110 | 7.94 |
| | | 48 | 14 | 2 | | 3.77 |
| | Fusiform gyrus | 45 | -58 | -16 | 1232 | 7.44 |
| | | 30 | -40 | -19 | | 6.71 |
| | | 42 | -43 | -22 | | 6.49 |
| | Fusiform gyrus | -42 | -70 | -10 | 522 | 6.76 |
| | | -45 | -55 | -13 | | 6.35 |
| | | -42 | -46 | -19 | | 6.29 |
| | Precentral gyrus | 33 | 5 | 62 | 72 | 5.27 |
| | | 36 | 17 | 59 | | 3.47 |
| | Inferior frontal gyrus | -48 | 35 | 23 | 67 | 5.25 |

| | | | | | | |
|---|---|---|---|---|---|---|
| | (lIFG) | -42 | 26 | 23 | | 4.69 |
| | Insula (lINS) | -30 | 26 | -7 | 132 | 5.21 |
| | Precentral gyrus | -42 | 5 | 29 | 86 | 4.58 |
| | | -48 | 11 | 26 | | 4.24 |
| | Caudate | -9 | 11 | -1 | 61 | 4.25 |
| | Intraparietal sulcus | -39 | -46 | 44 | 56 | 4.12 |
| | | -48 | -40 | 44 | | 3.83 |
| ***Interaction (LP>GP – GP>LP)*** | Paracentral lobe | 0 | -25 | 74 | 59 | 4.34 |
| | | 3 | -25 | 65 | | 4.16 |
| | | -9 | -22 | 71 | | 3.94 |

| **Cue-evoked** | **Regions** | **MNI coordinates** | | | **Cluster size** | **T** |
|---|---|---|---|---|---|---|
| | | **x** | **y** | **z** | | |
| ***Gain>Loss*** | Cingulate gyrus | -6 | -10 | 38 | 1108 | 5.38 |
| | | -9 | -1 | 44 | | 5.12 |
| | | -33 | -10 | 41 | | 5.11 |
| | Middle temporal gyrus | 42 | -64 | 8 | 252 | 5.25 |
| | | 51 | -52 | 11 | | 4.92 |
| | | 63 | -46 | 11 | | 4.88 |
| | Inferior occipital gyrus | 27 | -70 | -10 | 296 | 5.09 |
| | | 24 | -82 | 14 | | 4.55 |
| | | 15 | -88 | 11 | | 4.54 |
| | Middle temporal gyrus | -54 | -64 | 14 | 58 | 4.96 |
| | | -45 | -64 | 23 | | 4.36 |
| | Superior temporal gyrus | 51 | -4 | -13 | 47 | 4.84 |
| | | 60 | 2 | -13 | | 4.15 |
| | | 63 | -7 | -10 | | 3.88 |
| | Superior occipital gyrus | -15 | -88 | 11 | 82 | 4.31 |
| | | -6 | -88 | 8 | | 4.10 |
| | | -18 | -91 | 2 | | 4.04 |
| | Anterior cingulate cortex | -9 | 50 | 2 | 80 | 4.28 |
| | | 18 | 53 | 2 | | 4.03 |
| | | 6 | 50 | -1 | | 3.95 |

| **Conjunction** | **Regions** | **MNI coordinates** | | | **Cluster size** | **T** |
|---|---|---|---|---|---|---|
| | | **x** | **y** | **z** | | |
| ***Cue-evoked: G>L + Positive confidence effect*** | Superior frontal gyrus | -6 | 59 | 8 | 105 | 4.29 |
| | Orbital frontal cortex | -6 | 53 | -4 | | 3.94 |
| | Orbital frontal cortex | 6 | 53 | -4 | | 3.54 |

**Table S7. fMRI model-free analysis – GLM1**

Note: Whole-brain cluster-defining height threshold at uncorrected p <.001, k = 47. With the same threshold, no brain region survives in the contrast of [gain>loss], [complete>partial] nor the contrast of interaction. The regions highlighted with red are ROIs used in the main paper.

MNI: Montreal Neurological Institute; Qc: chosen option value; Qu: unchosen option value;

| Parametric effect | Regions | MNI coordinates x | y | z | Cluster size | T |
|---|---|---|---|---|---|---|
| ***Qc*** (***Positive effect***) | VMPFC | 6 | 38 | -4 | 107 | 4.25 |
| | | -3 | 41 | -1 | | 4.2 |
| | | 9 | 47 | -7 | | 4 |
| ***Qc*** ***(Negative effect)*** | DMPFC | 0 | 26 | 44 | 253 | 6.62 |
| | | -3 | 35 | 32 | | 4.06 |
| | Fusiform gyrus | 45 | -58 | -10 | 262 | 5.9 |
| | | 42 | -52 | -16 | | 5.86 |
| | | 36 | -40 | -25 | | 4.34 |
| | Occipital gyrus | -45 | -58 | -13 | 361 | 5.44 |
| | | -42 | -70 | -10 | | 5.39 |
| | | -36 | -82 | -10 | | 5.3 |
| | IFG | 45 | 14 | 29 | 262 | 5.12 |
| | | 42 | 5 | 26 | | 4.75 |
| | | 48 | 32 | 20 | | 4.5 |
| | Thalamus | 9 | -13 | 8 | 64 | 4.9 |
| | | -6 | -16 | 11 | | 3.53 |
| | Striatum | 21 | 8 | -7 | 64 | 4.68 |
| | | 15 | 2 | -19 | | 4.59 |
| | | 18 | 17 | -4 | | 4.01 |
| | Superior parietal lobe | 27 | -64 | 47 | 276 | 4.54 |
| | | 39 | -49 | 56 | | 4.49 |
| | | 24 | -58 | 35 | | 4.46 |
| ***Qu*** (***Positive effect***) | n.s. | | | | | |
| ***Qu*** (***Negative effect***) | Occipital gyrus | 24 | -85 | 17 | 119 | 4.67 |
| | | 12 | -94 | 14 | | 4.5 |
| | | 15 | -88 | 5 | | 4.17 |
| ***Context value: V*** (***Positive effect***) | Lingual gyrus | 24 | -64 | -7 | 67 | 4.65 |
| | | 27 | -55 | -10 | | 4.64 |
| | | 24 | -55 | -19 | | 3.82 |
| | Calcarine | -27 | -61 | 11 | 71 | 4.19 |
| | | -30 | -79 | 11 | | 4.1 |
| | | -24 | -91 | 17 | | 3.98 |
| | Occipital gyrus | 39 | -76 | 14 | 66 | 3.98 |
| | | 27 | -82 | 14 | | 3.84 |
| | | 18 | -85 | 14 | | 3.8 |
| ***Context value: V*** (***Negative effect***) | n.s. | | | | | |

**Table S8. fMRI model-based analysis – GLM3**
Note: Whole-brain cluster-defining height threshold at uncorrected p <.001, k = 47. The regions highlighted with red are ROIs used in the main paper.
MNI: Montreal Neurological Institute

| Parametric effect | Regions | MNI coordinates | | | Cluster size | T |
|---|---|---|---|---|---|---|
| | | x | y | z | | |
| ***Qc*** (***Positive effect***) | PCC | -9 | -13 | 41 | 1022 | 5.97 |
| | | -39 | -34 | 23 | | 5.29 |
| | | 30 | 20 | 17 | | 5.19 |
| | Cerebelum | 6 | -70 | -13 | 574 | 5.66 |
| | | 9 | -79 | -1 | | 5.23 |
| | | 21 | -85 | 5 | | 5.03 |
| | Occipital gyrus | -15 | -88 | 5 | 86 | 5.31 |
| | | -6 | -88 | 8 | | 4.49 |
| | Fusiform | -30 | -34 | -19 | 60 | 4.83 |
| | | -36 | -22 | -16 | | 4.61 |
| | | -42 | -25 | -7 | | 4.22 |
| | VMPFC | -3 | 53 | 2 | 105 | 4.16 |
| | | 0 | 65 | 11 | | 4.08 |
| | | 9 | 50 | 8 | | 3.83 |
| ***Qc*** (***Negative effect***) | n.s. | | | | | |
| ***Difference: \|Qc-Qu\| (Positive effect)*** | n.s. | | | | | |
| ***Difference: \|Qc-Qu\| (Negative effect)*** | Inferior temporal gyrus | 48 | -52 | -13 | 71 | 5.18 |
| | | 42 | -58 | -13 | | 4.97 |
| | | 39 | -40 | -16 | | 3.83 |
| | Occipital gyrus | -36 | -88 | -7 | 66 | 4.61 |
| | | -42 | -61 | -13 | | 4.49 |
| | | -39 | -73 | -10 | | 4.11 |
| ***Confidence$_{t-1}$ (Positive effect)*** | Insula | 42 | -19 | 5 | 83 | 4.99 |
| | | 36 | -22 | 17 | | 4.48 |
| | | 51 | -10 | 5 | | 4.13 |
| ***Confidence$_{t-1}$ (Negative effect)*** | Cerebelum | 36 | -49 | -28 | 15452 | 9.08 |
| | | 36 | -61 | -13 | | 8.75 |
| | | 24 | -64 | -19 | | 8.4 |
| | Cerebelum | -9 | -31 | -34 | 94 | 4.94 |
| | | 3 | -37 | -34 | | 4.8 |
| | | -9 | -31 | -25 | | 4.54 |
| | IFG | -45 | 44 | 8 | 71 | 4.73 |
| | | -48 | 38 | 14 | | 4.5 |

**Table S9. fMRI model-based analysis – GLM4**
Note: Whole-brain cluster-defining height threshold at uncorrected p <.001, k = 47.
MNI: Montreal Neurological Institute

#### Data & Code availability

Anonymized behavioral data, analyses scripts and second-level neuroimaging maps will be made available on public repositories upon acceptance of this manuscript.
